# Soil-microbes-mediated invasional meltdown in plants

**DOI:** 10.1101/2020.03.11.987867

**Authors:** Zhijie Zhang, Yanjie Liu, Caroline Brunel, Mark van Kleunen

## Abstract

While most alien species fail to establish, some invade native communities and become widespread. Many of these communities have been invaded by multiple aliens, suggesting that aliens may cause invasional meltdowns. Here, we tested whether and how a third plant species affects the competitive outcome between alien and native plants through its soil legacy. We first conditioned soil with one of ten species (six natives and four aliens) or without plants. Then, we grew on these 11 soils, five aliens and five natives without competition, and with intra- or interspecific competition (all pairwise alien-native combinations). We found that aliens were not more competitive than natives when grown on soil conditioned by other natives or on non-conditioned soil. However, aliens were more competitive than natives on soil conditioned by other aliens. Although soil conditioning rarely affected the strength of competition between later plants, soil conditioned by aliens changed the competitive outcomes by affecting growth of aliens less negatively than that of natives. Microbiome analysis confirmed this finding by showing that the soil-legacy effects of one species on later species were less negative when their fungal endophyte communities were less similar; and that fungal endophyte communities were less similar between two aliens than between aliens and natives. Our study suggests that coexistence between aliens and natives is less likely with more alien species. Such invasional meltdown is likely mediated by spill-over of fungal endophytes, some of which are pathogenic.

## Introduction

What determines invasion success of alien species is a central and urgent question in ecology^1^. Charles Elton, in his famous book, posited superior competitive ability as one of the mechanisms^2^. Since then, hundreds of experiments have studied competition between native and alien species, confirming that many successful alien species are indeed more competitive than natives^3–5^. Most studies, however, focused on pairwise interactions^6^ (i.e. between an alien and a native species; but see ref^7–9^ for studies of multispecies interactions), although in nature most species interact with multiple species. Moreover, interactions between alien species have also been frequently observed^10^. In many cases, aliens appear to favor other aliens over natives^11,12^, a phenomenon called invasional meltdown^13^. Still, invasional meltdown has so far mainly been studied in pairs of alien species without considering interactions with native species^14,15^. Therefore, the competitive outcome between alien and native species in multispecies communities remains unknown.

A major challenge in community ecology is to predict competitive outcomes in multispecies communities (e.g. to predict which species will dominate). Many studies suggest that outcomes in multispecies communities could be predicted from two-species systems, by assuming that interactions remain pairwise in all systems^16–18^, which. For a hypothetical example, consider adding a third species into a two-species community (Fig. 1b). If we know from previous pairwise experiments that the third species strongly suppresses one of the two species, we would predict that it will indirectly release the other species from competition. Although this ‘bottom-up’ approach is supported by several experiments on microbes^19,20^, the effect of one competitor on another (i.e. the strength of competition) can be changed by a third species^21,22^ (Fig. 1c & d). For example, it was shown that *Skeletonema costatum*, a cosmopolitan diatom, does not directly affect the growth of *Karenia brevis*, a dominant dinoflagellate in the Gulf of Mexico, but undermines the allelopathic effects of *K. brevis*^23^. This would also lessen the effect of *K. brevis* on other phytoplankton species, and interactions might consequently not always remain pairwise. Therefore, we need to test how the competitive outcome between alien and native species is affected by other species explicitly.

**Figure 1.**
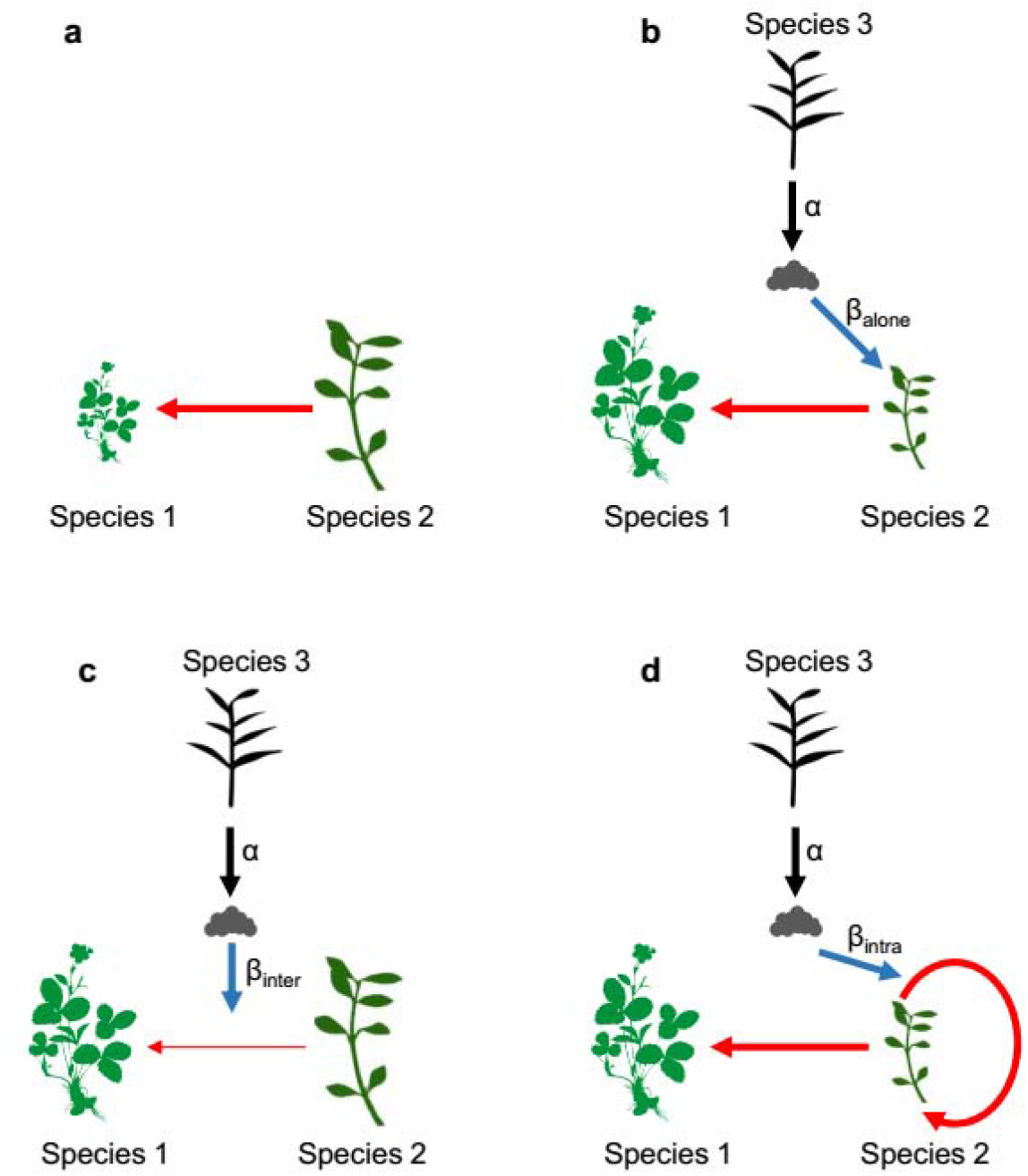
Graphical illustration of how a third species can affect competitive outcomes between two species through changes in soil microbial communities. **a**, In pairwise competition, species 2 suppresses species 1. Consequently, species 2 is more competitive, as indicated by its larger size. **b**, By modifying soil microbial communities (α; black arrow), species 3 favors species 1 by suppressing species 2 (β_alone_). Now species 1 is more competitive, as indicated by its larger size. **c**, Species 3 does not suppress species 2, but favors species 1 by lessening the suppression of species 2 on species 1 (β_inter_; indicated by the thinner red arrow). **d**, Species 3 favors species 1 by increasing the suppression of species 2 on itself (β_intra_; indicated by the presence of a curved red arrow). The overall effect of the third species on competitive outcomes between species 1 and 2, β_total_, is the net effect of β_alone_, β_inter_ and β_intra_.

Competition occurs through different processes, which makes it challenging to study. The most widely studied process is resource competition^24^, partly because competition for space, food and other resources is the most intuitive. Nevertheless, growing evidence shows that resource use alone cannot always explain success of alien species^25–27^. Competition can also act through other trophic levels. This so-called apparent competition^28^ has been extensively studied in systems in which plants affect others through shared aboveground herbivores^6,7^. The last two decades, however, has also seen an increased interest in apparent competition mediated by soil microbes. More and more studies reveal that plants modify soil microbes with consequences for their own development, and affecting plants that grow subsequently on the soil^29–33^ (a mechanism that we hereafter refer to as a soil-legacy effect).

Studies on soil-legacy effects have opened up new avenues to test mechanisms of plant invasion^34^, such as enemy release^35–37^ and novelty of aliens^38,39^. Based on these mechanisms, we would expect that the origin (alien or native) of the third species matters in how they affect competitive outcomes between alien and native species. First, enemy release posits that alien plants are released from their enemies^40^, and therefore soil conditioned by alien plants should accumulate few soil pathogens. Consequently, aliens would free natives that grow later on that soil from pathogens, unless they accumulate pathogens that are highly toxic to natives^41^. However, if aliens grow later on the soil, they might not be affected because they are already released from pathogens. Following this logic, soil conditioned by aliens would subsequently favor natives over aliens. Second, natives are familiar to (i.e. co-evolve with) each other, whereas aliens and natives are novel to each other (i.e. lack of coevolution)^39,42^. Therefore, natives should accumulate soil pathogens that are more likely shared with other natives than with aliens. Following this logic, soil conditioned by natives would subsequently favor aliens over natives. Whether these two expectations hold remains unknown.

To date, the competitive outcome between alien and native plants in multispecies communities remains unclear. We tested this with a large multi-species experiment. We first conditioned soil with one of ten species (six natives and four aliens) or, as a control, without plants. Then, on each of these 11 soils, five alien and five native test species were grown without competition, and with intraspecific or interspecific competition, using all pairwise alien-native combinations. To assess the potential role of microbes, we also analyzed the relationship between soil communities and soil-legacy effects. We addressed the following questions: (1) Does a third species (i.e. a soil-conditioning species) affect the competitive outcome between subsequent alien and native test species through a soil-legacy effect (β_total_, the net effect of β_alone_, β_inter_ and β_intra_ in Fig. 1), and does the origin (native or alien) of the third species matter? (2) If so, does the third species affect competitive outcomes through its soil-legacy effect on the growth of test species (β_alone_), or through its soil-legacy effect on the strength of competition (β_inter_ or β_intra_)? (3) Does variation in soil microbial communities among the conditioned soils explain the variation in soil-legacy effects ?

## Materials and Methods

### Study location and species

We conducted our experiment in the Botanical Garden of the University of Konstanz, Germany (47.69°N, 9.18°E). We conditioned soil with one of four plant species that are naturalized aliens in Germany and six plant species that are native to Germany. For these 10 soil-conditioning species, we tested their soil-legacy effects on five naturalized alien and five native species (test species; Table S1). The soil-conditioning and test species partly overlapped, and in total we used seven alien and six native species. We used multiple species to increase our ability to generalize the results^43^. The classification of the species as natives or naturalized aliens in Germany was based on the FloraWeb database^44^. Among the seven alien species, three are native to North America, one to Southern Africa, and three to other parts of Europe (Table S1). All 13 species can be locally abundant and are widespread in Germany (i.e. occur in at least 30% regions in Germany, see Table S1 for details). As widespread species are likely to have high spread rates, the alien species can be considered as invasive or probably invasive *sensu* Richardson, et al. ^45^. All species mainly occur in grasslands and overlap in their distributions according to FloraWeb, and thus are very likely to co-occur in nature.

Seeds of the native species and one of the alien species (*Onobrychis viciifolia*) were purchased from Rieger-Hofmann GmbH (Blaufelden-Raboldshausen, Germany). Seeds of the other species were from the seed collection of the Botanical Garden of the University of Konstanz. We initially planned to use the same species in the soil-conditioning and test phases. However, in the soil-conditioning phase, seeds of one of the six native species (*Cynosurus cristatus*) were contaminated with other species, and germination success of two aliens (*Solidago gigantea* and *Salvia verticillata*) was low. Therefore, we replaced these three species in the test phase with three alien species (*Solidago canadensis*, *Senecio inaequidens* and *Epilobium ciliatum*).

### Experimental setup

#### Soil-conditioning phase

From 18 June to 2 July 2018 (Table S1), we sowed the four alien and six native soil-conditioning species separately into trays (10 cm × 10 cm × 5 cm) filled with potting soil (Topferde^@^, Einheitserde Co., Germany). Seeds were not sterilized. Because we wanted the different species to be in similar developmental stages at the beginning of the experiment, we sowed the species at different times (Table S1), according to their germination timing known from previous experiments. We placed the trays in a greenhouse under natural light conditions, with a temperature between 18 and 25°C.

For each species, we transplanted 135 seedlings individually into 1.5-L pots. This was done for eight out of ten species, and done from 9 to 11 July 2018. For the other two species, *Sa. verticillata* and *So. gigantea*, we transplanted 61 and 115 seedlings respectively, from 25 July to 12 August (Table S1). This was because these two species germinated more slowly and irregularly than foreseen. We also added 330 pots that did not contain plants as a control treatment. In a complete design, we would have had 1680 pots. However, because we had fewer pots of *C. cristatus*. *So. gigantea* and *Sa. verticillata*, we ended up with 1521 pots. The substrate that we used was a mixture of 37.5% (v/v) sand, 37.5% vermiculite and 25% field soil. The field soil served as inoculum to provide a live soil microbial community, and was collected from a grassland site in the Botanical Garden of the University of Konstanz on 12 June 2018. We removed plant materials and large stones by sieving the field soil through a 1-cm mesh, and immediately thereafter stored it at 4°C until the transplanting.

After the transplanting, we randomly assigned the pots to positions in four greenhouse compartments (23°C/18°C day/night temperature, no additional light). Each pot sat on its own plastic dish to preserve water and to avoid cross-contamination through soil solutions leaking from the bottoms of the pots. Seedlings that died within two weeks after transplanting were replaced by new ones. All pots, including both the ones with and without plants, were watered as needed, randomized twice across the four compartments, and fertilized seven times during the soil-conditioning phase with an NPK water-soluble fertilizer (Universol Blue^®^, Everris, Germany) at a concentration of 1‰ m/v. From 22 to 26 October 2018, 15 weeks after the start of soil-conditioning phase, we harvested all soil. We cut aboveground biomass at soil level and freed the soil from roots by sieving it through a 5-mm mesh. The mesh was sterilized in between using 70% ethanol. For the pots in the control treatment, the soil was also sieved through the mesh. Then, we put the sieved soil of each pot separately into new 1-L pots, which were used in the test phase. So, as recommended by Brinkman et al. (2010)^46^, we did not pool soil from different pots in order to ensure independence of replicates. The collected aboveground biomass was dried at 70°C to constant weight, and weighed to the nearest 1 mg.

#### Test phase

From 9 to 18 October 2018, we sowed the five alien and five native test species (Table S1) in a similar way as we had done for the species of the soil-conditioning phase. On 29 and 30 October, we transplanted the seedlings into the 1-L pots filled with soil from the soil-conditioning phase. Three competition treatments were imposed (Fig. 2): 1) no competition, in which individuals were grown alone; 2) intraspecific competition, in which two individuals of the same species were grown together; 3) interspecific competition, in which one individual of an alien and one individual of a native species were grown together. We grew all ten species without competition, in intraspecific competition, and in all 25 possible native-alien combinations of interspecific competition. For the plants that were grown in non-conditioned soil, we replicated each species without competition 12 times, and with intraspecific competition and each interspecific native-alien combination six times. For the plants that were grown on conditioned soil, we had three replicates for each combination of a competition treatment (10 without competition, 10 with intraspecific competition, 25 with interspecific competition) and soil-conditioning species (six native and four alien). Because we had fewer replicates for soil conditioned with *C. cristatus*, *So. gigantea* and *Sa. verticillata*, the final design had 1521 pots (and 2639 individuals) in the test phase.

**Figure 2.**
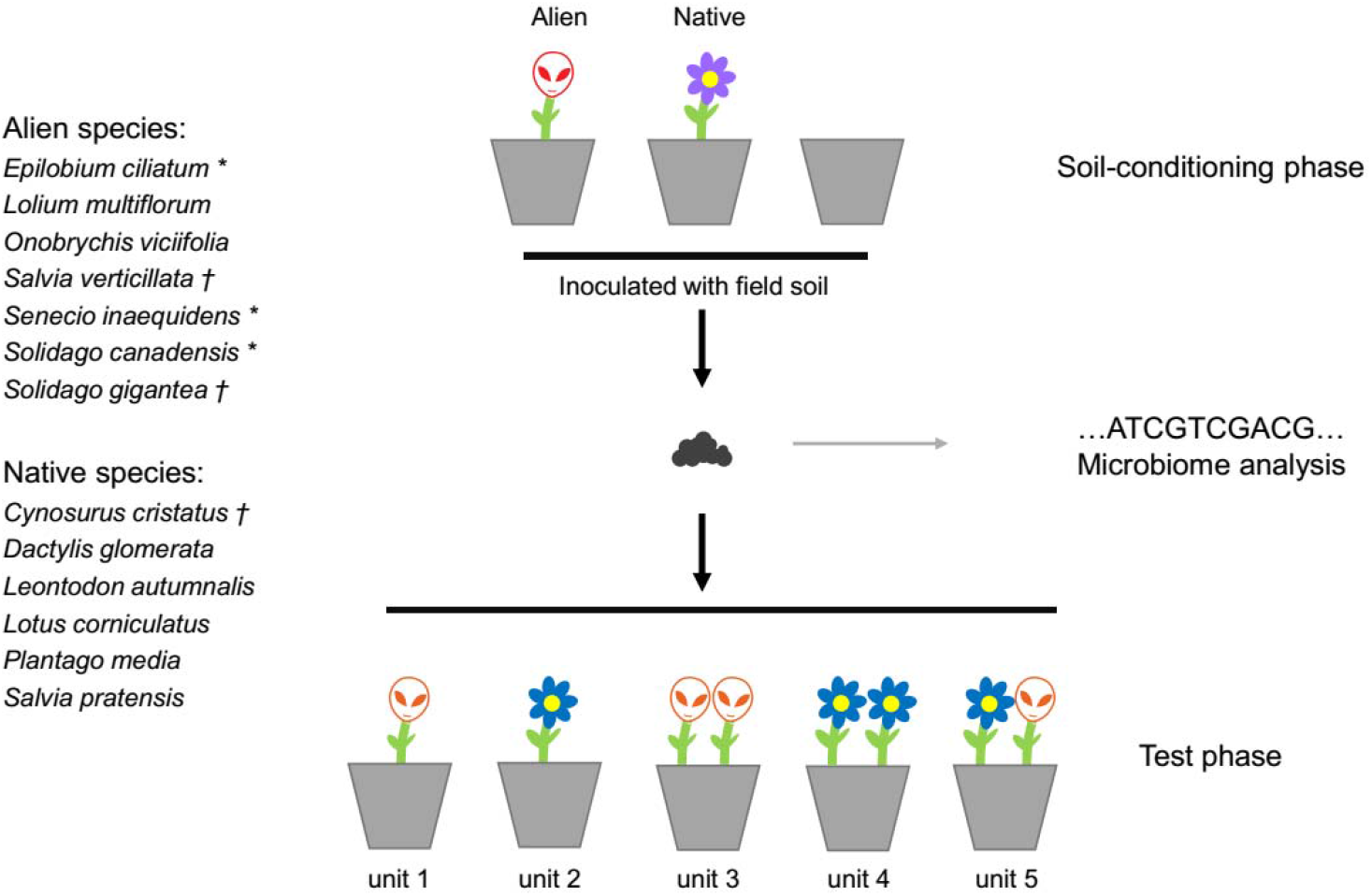
Graphical illustration of the experimental design. In the soil-conditioning phase, soil was conditioned by one of ten species (either alien or natives), or not conditioned. Then, test species were grown on each of these 11 soils alone or with intra- or interspecific competition. Soil was sampled after conditioning, and amplicon sequencing was used to assess the microbial communities. Plants grown alone (units 1-2) were used to test how soil-conditioning species affected the growth of test species (β_alone_ in Fig. 1). The differences between plants grown in competition (units 3-5) and the ones grown alone were used to test how soil-conditioning species affected the strength of intra- and interspecific competition (β_intra_ & β_inter_). Aboveground biomass across competition treatments indicated competitive outcomes (i.e. aliens are considered more competitive than natives when they had a higher aboveground biomass across units 1-5), and were used to test how soil-conditioning species affected competitive outcomes (β_total_). Species marked with asterisks were only used in the test phase. Species with daggers were only used in the soil-conditioning phase. Others were used in both phases.

We randomly assigned the pots to positions in three greenhouse compartments. Each pot sat on its own plastic dish. Seedlings that died within two weeks after transplanting were replaced with new ones. All plants were watered as needed, and fertilized four times during the test phase with the same fertilizer as that in the soil-conditioning phase. The pots were re-randomized across the three compartments on 10 December 2018. On 8 and 9 January 2019, ten weeks after the transplanting, we harvested all aboveground biomass of each plant. For the plants that were grown alone, we washed the belowground biomass free from soil. This could not be done for the plants with competition, as their roots were so tangled that we could not separate them. The biomass was dried at 70°C to constant weight, and weighed to the nearest 1 mg.

#### Soil sampling, DNA extraction, amplicon sequencing and bioinformatics

From 22 to 26 October 2018, when we harvested the soil from the soil-conditioning phase, we randomly selected six pots of each of the ten soil-conditioning species. For each of these pots, we homogenized the soil and then put a random sample of 10-20 ml in sterile plastic tubes (50 ml). We additionally collected soil from six of the pots without plants. The 66 samples were immediately stored at −80 °C until DNA extraction.

We extracted DNA from 0.25 g of each soil sample using the DNeasy® PowerSoil® Kit (Qiagen, Hilden, Germany), following the manufacturer’s protocol. PCR amplifications and amplicon sequencing were then performed by Novogene (Beijing China). The V3-V4 region of bacterial 16S rDNA gene was amplified in triplicate with the universal primers 341F/806R (forward primer: 5’-CCTAYGGGRBGCASCAG-3’; reverse primer: 5’-GGACTACNNGGGTATCTAAT-3’^47^). The ITS2 region of the fungal rDNA gene was amplified in triplicate with the primers specific to this locus (forward primer: 5’-GCATCGATGAAGAACGCAGC-3’; reverse primer: 5’-TCCTCCGCTTATTGATATGC-3’^48^).

We processed the raw sequences with the *DADA2* pipeline, which was designed to resolve exact biological sequences (Amplicon Sequence Variants). After demultiplexing, we removed the primers and adapter with the *cutadapt* package^49^. We trimmed the 16S sequences to uniform lengths. Sequences were then dereplicated, and the unique sequence pairs were denoised using the *dada2* package^50^.We then merged paired-end sequences, and removed chimeras. We rarefied bacteria and fungi to 30,000 and 9,500 reads, respectively, to account for differences in sequencing depth. Three samples with lower reads for bacteria or fungi, and two samples with low amplicon concentrations for fungi were excluded from analyses. For fungi, we assigned the sequences to taxonomic groups using the UNITE^51^ database. Then, we identified putative fungal functional groups that could affect plant fitness using the FUNGuild database^52^. Sequence variants assigned to arbuscular mycorrhizal fungi, plant pathogens and endophytes represented respectively <0.1%, 11.4% and 15.7% of the total read abundance. Sixty-five sequence variants were assigned as both pathogens and endophytes, representing 6.3% of the total read abundance. This indicates that *c.* 40% of the assigned endophytes are pathogenic. Because assigned arbuscular mycorrhizal fungi had extremely low abundance and were not detected in 37 out of 62 soil samples, we did not analyze the data of arbuscular mycorrhizal fungi.

### Statistical analyses

All statistical analyses were done in R, version 3.6.1^53^. We provide the main information for each model in the main text, and details (e.g. random effects, variance structure) in Supplement S2.

#### Analyses of plant performance

To test whether soil-conditioning plants affected competitive outcomes between alien and native species (β_total_, the net effect of β_alone,_ β_intra_ and β_inter_) and the strength of competition (β_inter_ in Fig. 1c and β_inter_ in Fig. 1d) in the test phase, we used a linear mixed-effect model (Model.plant.1), as implemented in the *nlme*^54^ package. The model included aboveground biomass of the test plants as the response variable, and the soil-conditioning treatment (none, same species as the test species, native species, alien species), competition treatment (no, intra- and interspecific competition), origin of test species (native, alien) and their interactions as the fixed effects. A significant interaction between competition treatment and soil-conditioning treatment would indicate that soil-conditioning treatments affects the strength of competition. A significant three-way interaction of competition treatment, soil-conditioning treatment and origin of the test species would indicate that the soil-conditioning treatments affect the strength of competition of alien and native plants differently. A significant interaction between soil-conditioning treatment and origin of the test species would indicate that the soil-conditioning treatment affects biomass production of alien and native test species differently, averaged across all competition treatments. In other words, it would indicate that the soil-conditioning treatment affects the competitive outcome between aliens and natives. Competitive outcome here refers to which species will exclude or dominate over the other species at the end point for the community^55^. Most studies infer the competitive outcome by only growing the species in mixture^56^. However, we inferred it from the average of plants without competition, in monocultures and in mixtures. This method has the advantages that it better mimics the dynamics of species populations across space and time^5,57^, and that it increases the precision of estimating competitive outcomes^55^.

To test whether soil-conditioning plants directly affected growth of alien and native species (β_alone_ in Fig. 1b), we analyzed the subset of test plants grown without competition with linear mixed-effect models (Model.plant.2). These models included aboveground, belowground or total biomass of test plants as the response variables, and soil-conditioning treatment, origin of the test species and their interaction as fixed effects. For all mixed-effect models, the significance of fixed effect was assessed with likelihood-ratio tests when comparing models with and without the effect of interest^58^.

The soil-conditioning treatment had four levels: 1) the soil was not conditioned by any plant (non-conditioned soil), 2) the soil was conditioned by the same species as the focal test plant (home soil), and if the soil was conditioned by another species, this was either 3) an alien species (alien soil) or 4) a native species (native soil). We created three dummy variables^59^ to split up these four soil-conditioning treatments into three contrasts to test: 1) Does it matter whether the soil was conditioned by plants or not (Soil_Non-conditioned/Conditioned_)? 2) When the soil was conditioned by plants, does it matter whether the soil was conditioned by the same species as the focal test plant or by a different species (Soil_Home/Away_)? 3) When the soil was conditioned by a species different from the focal test plant, does it matter whether the soil was conditioned by an alien or a native species (Soil_Alien/Native_)?

Likewise, for the first model (Model.plant.1), which used data from all competition treatments, we created two dummy variables to split up the three competition treatments – no, intra- and interspecific competition – into two contrasts to test: 1) Does it matter whether the test plant was grown with competition (Comp_Yes/No_)? 2) When the test plant was grown with competition, does it matter whether the competitor belonged to the same species or not (Comp_Intra/Inter_)?

In a few cases of the interspecific competition treatment (103 out of 1573 pots), competitor species were the same as the soil-conditioning species. Therefore, these pots are testing a two-species rather than a three-species interaction. However, removing these data points does not affect the results, indicating that our results are robust (Table S2). It could be that soil-legacy effects are not due to differences in microbial communities of the soil but due to differences in nutrient availability^60^. For example, larger soil-conditioning plants may have left fewer nutrients in the soil, resulting in decreased growth of subsequent test species. To account for this, we added aboveground biomass of the soil-conditioning plant as the covariate in Model.plant.1. We found that aboveground biomass of test plants decreased with that of the soil-conditioning plant (Fig. S1), indicating that nutrient availability might affect test plants. However, adding the covariate did not affect the significance of the other effects (Table S3), indicating that our results are robust.

#### Analyses of the soil microbial community

To test the effect of soil-conditioning species on soil microbial communities (α in Fig. 1), we used three methods. First, we tested whether the presence of a soil-conditioning plant affected the composition of soil microbial communities, and whether this effect depended on the origin of the soil-conditioning species. To do so, we used permutational analysis of variance (PERMANOVA), as implemented in the *adonis* function of the *vegan* package^61^ (Model.soil.1). The models included reads relative abundances of bacteria or fungi as the response variables and soil-conditioning treatment as the explanatory variable. We split up the three soil-conditioning treatments into two contrasts to test: 1) Does it matter whether the soil was conditioned by plants or not? 2) When the soil was conditioned by plants, does it matter whether the species is alien or native?

Second, we tested whether alien and native species accumulated putative fungal pathogens, which were identified from FUNGuild, to different degrees. To do so, we used linear mixed models (Model.soil.2) that included the species richness, Shannon diversity or relative abundance of fungal pathogens as the response variable, and soil- conditioning treatments, which were again split up into two contrasts, as the fixed effect. Because some bacteria might be pathogenic, and 70% of the fungi could not assigned to functional groups based on FUNGuild, we also applied this analysis to species richness and Shannon diversity of all bacteria and fungi.

Third, we analyzed how conditioned soil communities differed 1) among plants from the same alien plant species, 2) among plants from the same native species, 3) among different alien species, 4) among different native species, and 5) between alien and native species. To do so, we used linear mixed models (Model.soil.3) and included averaged Bray-Curtis dissimilarities as the response variable, and the five above-mentioned categories of plant combinations as the fixed effect. The Bray-Curtis dissimilarities of bacteria, fungi, fungal pathogens or fungal endophytes were first calculated between all possible pairs of samples, and then averaged across replicates to get average values for each within-species pair or between-species pair. We split up the five categories of plant combinations into four contrasts to test: 1) Are soil communities more similar (or different) when conditioned by the same plant species than by another species? 2) When conditioned by the same species, are soil communities more similar for alien species than for native species? When conditioned by different species, 3) are soil communities more similar between two alien species than between an alien and a native species, and 4) are soil communities more similar for the latter than between two native species? We used heatmaps to visualize the community dissimilarities, whose values were mean-centered and then bounded to range from −1 to 1. This was done with the *corrplot* package^62^.

After testing the effect of soil-conditioning species on soil bacterial and fungal communities (α in Fig. 1), we aimed to identify which aspect of soil microbes explained the legacy effect of soil-conditioning species on test plants (i.e. which component of α explained the βs in Fig. 1). Because the analyses of plant performance revealed that the third species rarely significantly affected the strength of competition (i.e. on average, β_inter_ and β_intra_ did not differ significantly from 0), we present the analyses of effects of α on β_inter_ (or β_intra_) in the supplement S6.

We first tested whether diversity and abundance of potential soil enemies (one aspect of α) explained the soil-legacy effect on growth of test plants (β_alone_). To do so, we used linear mixed models (Model.link.1) and included the soil-legacy effect (β_alone_) as the response variable, and diversities of all soil bacteria, all fungi or the subset of fungal pathogens (or the relative abundance of fungal pathogens) as the fixed effects. Because the enemy release hypothesis predicts that alien species should have less chance to encounter enemies than native species^35,63^, we also added origin of test species and their interaction with diversities of soil bacteria, fungi or fungal pathogens (or relative abundance of fungal pathogens) as fixed effects. The soil-legacy effect, *β*_*alone,i,j*_ was calculated as

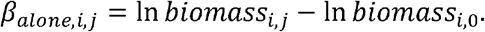

Here, ln *biomass_i,j_* and ln *biomass*_*i*,0_, are mean aboveground biomass of test species *i* when grown without competition (alone) on soil conditioned by species *j* and on soil not conditioned by plants, respectively. Positive values indicate that soil-conditioning species *j* improved growth of test species *i*.

Second, we tested whether microbial community dissimilarity (another aspect of α) between the soil-conditioning and test species explained the soil-legacy effect (β_alone_). To do so, we used linear mixed models (Model.link.2) and included the soil-legacy effect, *β*_*alone,i,j*_, as the response variable, and included Bray-Curtis dissimilarities between soil-conditioning and test species as the fixed effect. Because three out of ten test species were not included in the soil-conditioning phase, we could not calculate the microbial community dissimilarity between them and the soil-conditioning species. Consequently, this analysis was restricted to a subset (i.e. 70 out of 100 soil-conditioning species × test species pairs).

## Results

### Do soil-conditioning species affect differences in biomass production (i.e. competitive outcomes)?

On average, plants produced less aboveground biomass (−67.2%; χ^2^ = 10.31, *P* = 0.001) on conditioned soil than on non-conditioned soil, and on home soil (i.e. soil conditioned by the same plant species) than on away soil (−22.7%; χ^2^ = 4.54, *P* = 0.033; Fig. 3a; Table 1). Biomass of alien and native plants did not significantly differ across soil-conditioning treatments and competition treatments (χ^2^ = 0.083, *P* = 0.774; Fig. 3a; Table 1). Compared to non-conditioned soil, conditioned soil did not change the difference in biomass between alien and native plants across competition treatments (Origin × Soil_Non-conditioned/Conditioned_ interaction: χ^2^ = 1.395, *P* = 0.238). However, when grown on alien soil (i.e. soil conditioned by an alien plant), alien plants produced significantly more aboveground biomass (+18.2%) than native plants, whereas on native soil, this difference was smaller (+9.9%; Origin × Soil_Alien/Native_ interaction: χ^2^ = 4.74, *P* = 0.029; Fig 3a; Table 1). This indicates that soil conditioning with an alien plant pushed the competitive outcome more strongly towards subsequent aliens than soil conditioning with a native plant.

**Figure 3.**
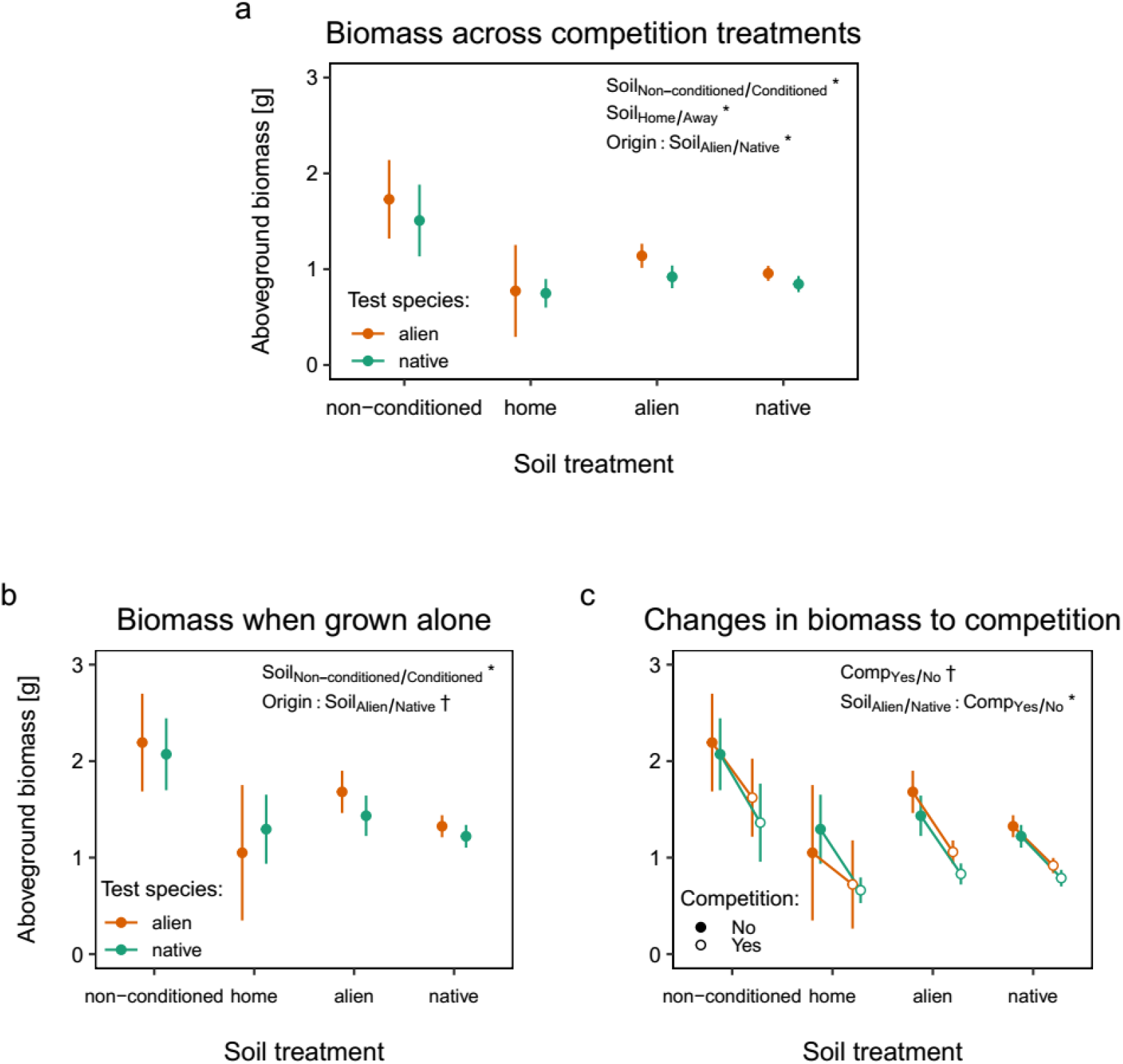
Effects of soil-conditioning treatments on aboveground biomass of alien (orange) and native (green) test species. **a**, Mean values (± SEs) were calculated across competition treatments. Alien test species are considered more competitive than natives when they had a higher aboveground biomass. **b**, Mean values were calculated based on aboveground biomass of plants grown alone. **c**, Slopes indicate the strength of competition, that is, the difference in aboveground biomass between plants grown alone (solid dots, the same values as in b) and in competition (open dots). For the soil-conditioning treatments, ‘non-conditioned’ refers to soil that was not conditioned by any plant, ‘home’ to soil conditioned by the same species as the test species, and ‘alien’ and ‘native’ to soils conditioned by other species than the test species, which were alien or native, respectively. Differences in mean values between different soil treatments in **a**, **b** and **c** indicate differences in β_total_, β_alone_ and β_inter_ (or β_intra_), respectively. See Fig. 1 for details on βs.

**Table 1.** Effects of soil treatments, competition treatments, origin of test species and their interactions on aboveground biomass of plants. Significant effects (P < 0.05) are in bold and marked with asterisks, and marginally significant effects (0.05 ≤ P < 0.1) are in italics and marked with a dagger symbol.

| | $\chi^2$ | **P** |
| --- | --- | --- |
| Transplanting date | 12.815 | **<0.001\*** |
| Soil<sub>Non-conditioned/Conditioned</sub> | 10.306 | **0.001\*** |
| Soil<sub>Home/Away</sub> | 4.535 | **0.033\*** |
| Soil<sub>Alien/Native</sub> | 0.107 | 0.744 |
| Origin (O) | 0.083 | 0.774 |
| Comp<sub>Yes/No</sub> | 3.738 | *0.053†* |
| Comp<sub>Intra/Inter</sub> | 0.795 | 0.373 |
| O : Soil<sub>Non-conditioned/Conditioned</sub> | 1.395 | 0.238 |
| O : Soil<sub>Home/Away</sub> | 1.669 | 0.196 |
| O : Soil<sub>Alien/Native</sub> | 4.741 | **0.029\*** |
| Soil<sub>Non-conditioned/Conditioned</sub> : Comp<sub>Yes/No</sub> | 0.956 | 0.328 |
| Soil<sub>Non-conditioned/Conditioned</sub> : Comp<sub>Intra/Inter</sub> | 0.176 | 0.675 |
| Soil<sub>Home/Away</sub> : Comp<sub>Yes/No</sub> | 0.121 | 0.728 |
| Soil<sub>Home/Away</sub> : Comp<sub>Intra/Inter</sub> | 2.273 | 0.132 |
| Soil<sub>Alien/Native</sub> : Comp<sub>Yes/No</sub> | 4.846 | **0.028\*** |
| Soil<sub>Alien/Native</sub> : Comp<sub>Intra/Inter</sub> | 0.321 | 0.571 |
| O:Comp<sub>Yes/No</sub> | 0.249 | 0.618 |
| O:Comp<sub>Intra/Inter</sub> | 0.371 | 0.542 |
| O : Soil<sub>Non-conditioned/Conditioned</sub> : Comp<sub>Yes/No</sub> | 0.511 | 0.475 |
| O : Soil<sub>Non-conditioned/Conditioned</sub> : Comp<sub>Intra/Inter</sub> | 0.001 | 0.972 |
| O : Soil<sub>Home/Away</sub> : Comp<sub>Yes/No</sub> | 1.725 | 0.189 |
| O : Soil<sub>Home/Away</sub> : Comp<sub>Intra/Inter</sub> | 0.156 | 0.693 |
| O : Soil<sub>Alien/Native</sub> : Comp<sub>Yes/No</sub> | 0.197 | 0.657 |
| O : Soil<sub>Alien/Native</sub> : Comp<sub>Intra/Inter</sub> | 0.000 | 0.990 |
| **Random effects** | **SD** |  |
| Family (focal test) | 0.165 |  |
| Species (focal test) | 0.199 |  |
| Family (competitor test) | 0.065 |  |
| Species (competitor test) | 0.076 |  |
| Family (soil) | 0.038 |  |
| Species (soil) | 0.031 |  |
| Residual | 0.187 |  |

### Do soil-conditioning species affect growth and the strength of competition?

For the subset of plants grown alone (competition-free), aboveground biomass was lower on conditioned soil than on non-conditioned soil (−59.8%; χ^2^ = 13.38, *P* < 0.001; Fig. 3b; Table S4). The competition-free plants also tended to produce less biomass on home soil than on away soil (Fig. 3b). This effect was not significant for aboveground biomass, but was marginally significant for belowground biomass (−15.0%; χ^2^ = 2.93, *P* = 0.087; Fig. 3b & S2; Table S4). Averaged across all soil-conditioning treatments, alien and native competition-free plants did not differ in biomass production (χ^2^ = 0.025, *P* = 0.875). However, aliens achieved more aboveground biomass (+17.3%) than natives on alien soil, whereas on native soil, this difference was smaller (+8.5%; Fig. 3b; Table S4). Although this difference was only marginally significant for aboveground biomass (χ^2^ = 2.90, *P* = 0.088) and belowground biomass (χ^2^ = 3.23, *P* = 0.072), it was significant for total biomass (χ^2^ = 4.56, *P* = 0.033; Table S4; Fig. S2). This result indicates that soil conditioning with an alien plant reduced growth of subsequent alien plants to a lesser degree than growth of subsequent native plants.

Competition reduced aboveground biomass (−35.1%; χ^2^ = 3.74, *P* = 0.053; Fig. 3c; Table 1), and was more intense when the test plants were grown on alien soil than on native soil (−39.3% *vs.* −33.0%; χ^2^ = 4.85, *P* = 0.028; Fig 3c; Table 1). However, the strength of competition was not affected by the other soil-conditioning treatments. Alien and native test species did not differ in their biomass responses to competition (χ^2^ = 0.25, *P* = 0.618), and this finding holds for each of the soil-conditioning treatments. We also found that intra- and interspecific competition did not differ in strength (χ^2^ = 0.80, *P* = 0.373), and that this finding holds for alien and native test species, and for each of the soil-conditioning treatments (Fig 3c; Table 1).

### Do soil microbial communities explain the soil-legacy effect?

Overall, the presence of plants significantly modified the composition of soil bacterial and fungal communities (Supplement S4.1). Moreover, alien and native plant species modified the composition of bacterial and fungal communities differently (Supplement S4.1). However, neither the presence of plants nor the origin of plants significantly affected relative abundance of fungal pathogens and diversities of bacteria, all fungi and the subset of fungal pathogens (Supplement S4.2). Further analyses showed that, the legacy effect of soil-conditioning species on test species that were grown alone (β_alone_) was not correlated to relative abundance of fungal pathogens and diversities of bacteria, fungi and fungal pathogens, and that this holds for both native and alien test species (Supplement S5.1).

The compositions of soil bacterial communities were less similar (i.e. more blue colors in Fig. 4) between individual plants of different species than between plants of the same species (χ^2^ = 4.31, *P* = 0.038; Fig. 4a & e; Table S9). Although this was not the case for fungal communities, their dissimilarity depended on the origins of the species in the between-species combination (Fig. 4b-d & f-h; Table S9). Specifically, compositions of fungal communities as a whole and of the subset of fungal endophytes were less similar between two alien plant species than between an alien and a native species (Fungi: χ^2^ = 4.00, *P* = 0.045; Fungal endophytes: χ^2^ = 12.11, *P* = 0.001). In addition, the compositions of fungal endophyte communities were less similar between an alien and a native species than between two natives (χ^2^ = 10.53, *P* = 0.001; Fig. 4d & h; Table S9).

**Figure 4.**
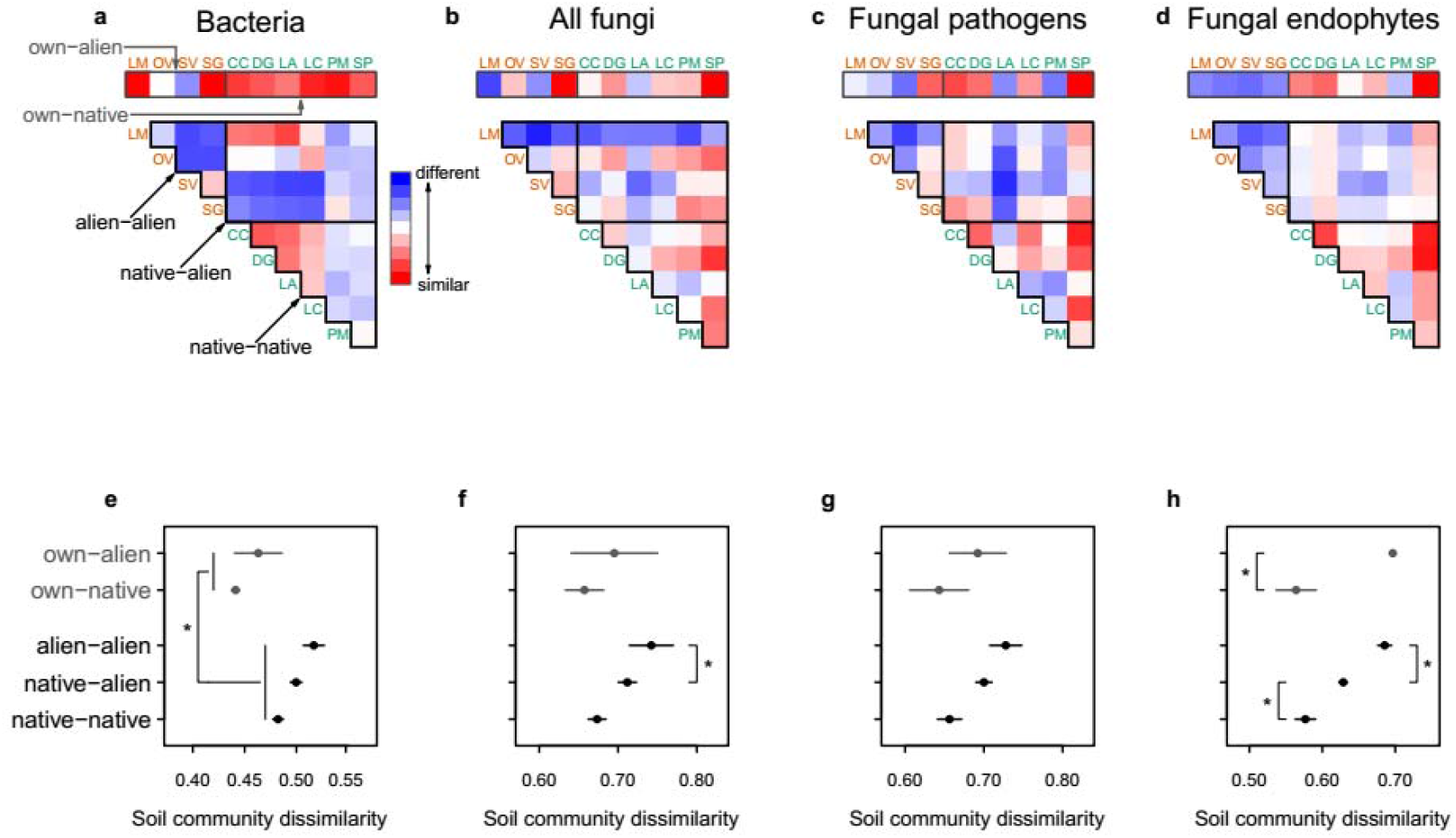
Dissimilarities of soil microbial communities within and between plant species. **a** & **e**, bacterial communities; **b** & **f**, fungal communities, **c** & **g**, fungal pathogen communities; **d** & **h**, fungal endophyte communities. The upper panels show the heatmaps of community dissimilarities of all within-species (top horizontal bars) and between-species combinations (triangular matrices), which are divided into five categories (own-alien, own-native, alien-alien, native-alien, native-native) with black borders. Labels at the top and along the diagonal provide abbreviations of species names (full names in Table S1) of aliens (orange) and natives (green). The colors in the heatmaps represent the relative dissimilarity, with the darkest blue hue representing the highest dissimilarity. The lower panel shows the mean values (±SEs) of each of the five categories. Significant differences between categories are indicated with an asterisk (i.e. α in Fig. 1 differs between categories). Own-alien: between individual plants of the same alien plant species; own-native: between plants of the same native species; alien-alien: between plants of different alien species; alien-native: between plants of alien and native species; native-native: between plants of different native species.

For the subset data on dissimilarities of soil communities between soil-conditioning and test species, we found that the legacy effect of soil-conditioning species on test species grown alone (β_alone_) became less negative with decreasing similarity of their fungal endophyte communities (χ^2^ = 7.49, *P* = 0.006; Fig. 5d; Table S13). A similar marginally significant trend was found for bacteria (χ^2^ = 2.78, *P* = 0.096; Fig. 5a; Table S13). For the other groups of microbiota, i.e. fungi overall and fungal pathogens, the soil-legacy effect (β_alone_) was not significantly correlated to the dissimilarity of soil communities (Fig. 5b & c; Table S13).

**Figure 5.**
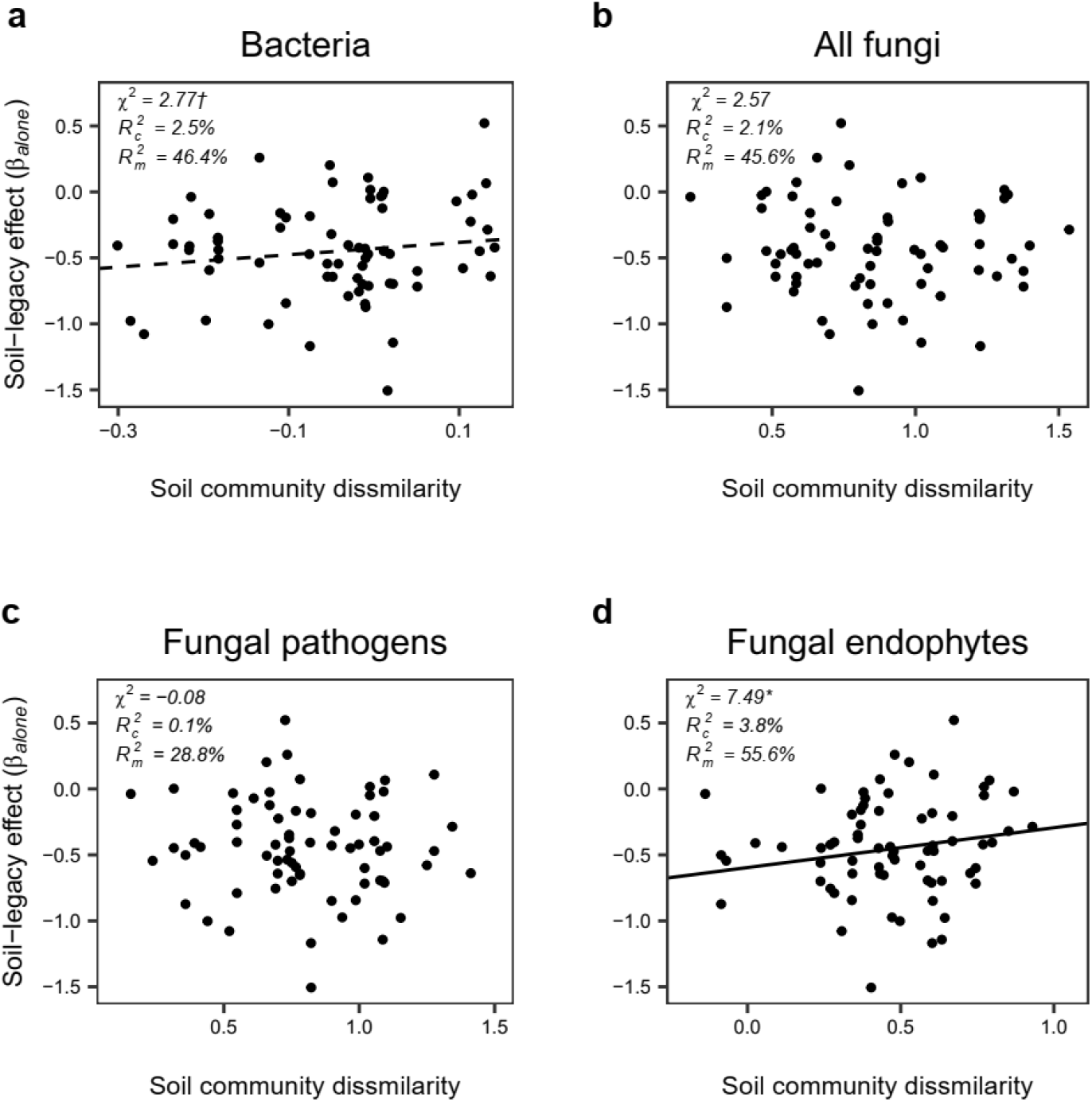
Effects of soil-community dissimilarity between soil-conditioning and test species on soil-legacy effects. **a**, bacterial communities; **b**, fungal communities; **c**, fungal pathogen communities; **d**, fungal endophyte communities. Negative values of the soil-legacy effect indicate that plants grew worse on conditioned soil than on non-conditioned soil. Soil-community dissimilarity was logit-transformed. Significant effects of community dissimilarity on soil-legacy effects are indicated with an asterisk (i.e. significant effect of α on β_alone_), and marginally significant effects with a dagger symbol. Chi-squared value (*χ*^2^), conditional R squared 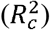 and marginal R squared 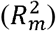 are reported in each panel.

## Discussion

We found that when grown on soil that had not been conditioned by plants, alien and native plants produced similar biomasses across competition treatments. The same was true on soil that had been conditioned by plants. This indicates that overall, the naturalized aliens in our study were not more competitive than natives, and that the presence of soil-conditioning species did not change this competitive outcome. However, on soil that had been conditioned by aliens, aliens produced more biomass than natives and thus were more competitive. Analysis of biomass of plants grown alone (without competition) indicated that conditioning by aliens changed the competitive outcomes by affecting growth of aliens less negatively than that of natives. The strength of competition, however, was rarely affected by the soil-conditioning treatment. Our analysis of soil microbiomes revealed that the legacy effect of soil-conditioning species on test species became less negative as their fungal endophyte communities became less similar, and that fungal endophyte communities were less similar between two aliens than between aliens and natives. This suggests that the less negative effect of conditioning by aliens on other aliens is partly driven by a lower chance of spill-over of pathogenic fungal endophytes between aliens.

### Invasional meltdown in a multispecies context

The similar aboveground biomass of aliens and natives on soil that had not been conditioned or had been conditioned by native plants indicates that on those soils aliens are not more competitive than natives. This result is in line with the recent finding that alien and native species do not differ in their competitive abilities if both of them are widespread and abundant species^5^, as was the case in our study. However, on soil conditioned by aliens, aliens were more competitive than natives. This finding supports the idea of invasional meltdown^3,13,14^ and partly explains the frequent co-occurrence of alien species^10^. So far, over 13,000 plant species have become naturalized outside their natural ranges^64,65^, and in some regions more than half of the flora consists of naturalized alien species^66^. These numbers are still increasing^67^, which means that interactions between alien species are likely to become more and more frequent. Our findings indicate that the relative facilitation between aliens, mediated by soil microbes, may accelerate the naturalization of aliens and their competitive impacts on natives.

Still, alien plants may not increase their abundance indefinitely, because intraspecific competition is generally stronger than interspecific competition^68^. We nevertheless did not find a difference between the strengths of intra- and interspecific competition. Probably, resource competition was not intense in our study as we fertilized the plants regularly. It is worth noting, however, that in our study, plants grew worse on home soil than on away soil. In other words, intraspecific apparent competition (soil-microbes-mediated intraspecific competition) was stronger than interspecific apparent competition. Consequently, alien plants were still self-limited. However, alien plants would gain an advantage if they were less limited by intraspecific apparent competition than natives were, which was supported by many studies^30^ but not ours. One possible reason for this discrepancy is the low statistical power in our study. Only two of the five alien test species were grown on home soil as we partly had different species in the soil-conditioning and test phases. Another reason could be that we used successful native species (i.e. widespread and locally abundant). Their intraspecific apparent competition might be weaker than for less successful native species^69^, and thus similar with that of the successful aliens.

It is debated in ecology whether it is possible to predict competitive outcomes in multispecies communities solely based on pairwise interactions. The results of our experiment suggest that this indeed is possible. For example, from the data of plants that were grown alone, which tested pairwise interactions between soil-conditioning and test species, we showed that alien test species produced more biomass than natives on soil that had been conditioned by aliens. This finding still holds when we also included the data of plants that were grown with competition to assess competitive outcomes in multispecies communities. Moreover, the soil-conditioning species rarely changed the strength of competition. When they did, they affected the strength of competition equally for alien and native species, and thus did not affect competitive outcomes. This finding echoes those of some other experiments. For example, a phytoplankton experiment by Prince, et al. ^23^ found that the strength of competition was modified only in two out of the ten species in their study. Friedman, et al. ^20^ found that competitive outcomes in three-species bacterial communities were predicted by pairwise outcomes with an accuracy of 90%. Therefore, we might in most cases be able to scale up from pairwise interactions to at least three-species interactions.

However, it might be too soon to scale up to systems with more than three species. Friedman, et al. ^20^ found that pairwise outcomes alone poorly predict outcomes of seven- or eight-species bacterial communities. This could indicate that with increasing diversity the likelihood increases that the strength of pairwise competition is modified by at least one of the many other species in the community. Future experiments that test competition between alien and native organisms in more diverse communities could shed light on this hypothesis. However, as competition occurs locally^70^, it is unlikely that more than a handful of species compete at the same time. Consequently, we believe that our experiment and results are representative for plant invasions in the real world.

### Potential mechanisms underlying invasional meltdown

We did not find evidence for release of soil enemies^71^. At the end of the soil-conditioning phase, alien and native plant species did not differ in the diversity and relative abundance of fungal pathogens. In addition, diversity and relative abundance of fungal pathogens in the soil did not significantly explain the performance of alien and native species in the test phase. This lack of evidence for enemy release contrasts with the findings that enemy release contributed to plant invasion^72,73^, but see ref^63^. This discrepancy may first arise from the incomplete information on the functional roles of bacteria and fungi. The functional roles of bacteria are hard to identify, and over 70% of the ITS reads in our study could not assigned to functional groups using FUNGuild. In addition, previous studies mainly focused on aboveground enemies and on herbivores. Belowground microbial enemies are more diverse and far less known, and many of them might be rare or less harmful. Therefore, diversity and relative abundance of soil pathogens may be less likely to capture the mechanism underlying soil-legacy effect than the actual identities of the pathogens. Indeed, we found that alien and native plants modified the composition of soil microbial communities in different ways (Supplement S4.1). This suggests that the soil-legacy effect is mainly mediated by the community structure of the soil microbial communities and less by the diversity and abundance.

Interestingly, we found that the compositions of fungal endophyte communities were less similar between alien plant species than between aliens and natives, and less similar than between natives. We found, however, a similar pattern when the field-soil inoculate used in the soil-conditioning phase had been sterilized (Fig. S8; Table S10). This suggests that the high dissimilarity of fungal endophyte communities between aliens is likely driven by endophytes that were already present in the plants before transplanting (e.g. as seed microbiota) rather than by those that were in the field-soil inoculum.

There are three potential reasons why compositions of fungal endophyte communities were less similar between two aliens than between other origin combinations. First, as we found that fungal endophyte communities became less similar with increasing phylogenetic distance between plant species (Supplement S7), it could be that the phylogenetic distance between aliens was higher than between aliens and natives, and also higher than between natives. However, as this was not the case (Supplement S7), this explanation can be discarded. A second potential explanation could be that natives have co-occurred with each other for a longer time, and thus share more similar endophytes^74^. A third potential explanation could be that if the alien species brought endophytes with them from their native ranges^75^, these endophytes jumped over to native hosts. Such host shifts of endophytes are more likely to involve native plants than other alien plants, as alien-native interactions are still more common than alien-alien interactions. Regardless of the exact reason, the observed differences in fungal endophyte communities suggest that they might play a role in the difference in soil-legacy effects.

In line with this idea, we found that the legacy effect of soil-conditioning species on test species became less negative with decreasing similarity in their fungal endophyte communities. As about 40% of the assigned endophytes were pathogenic, the overall effect of endophytes might be negative. Consequently, when one plant species cultivated very different endophyte communities compared to another, endophytes remaining in the soil matrix (e.g. root endophytes) were unlikely to infect and negatively affect the other. This finding, together with the higher difference in fungal endophyte communities between alien plant species than between alien and native plants species, provides a novel explanation for invasional meltdown. Still, the roles of endophytes are not well understood. Their effects depend on the environment and can range from pathogenic to mutualistic^76,77^. As a result, the legacy effect mediated by endophytes might even change with soil type. More experimental evidence for their role in soil-legacy effects and plant invasions is required. Manipulative studies based on synthetic microbial communities^78^ might shed light on the roles of endophytes in plant competition and invasion success.

## Conclusions

Our results indicate that the accumulation of alien species may be accelerated in the future, because aliens could favor other aliens over natives through soil-legacy effects, mediated by soil microbial communities (i.e. apparent competition). Since Charles Darwin^39^, novelty has been posited as an important mechanism of invasion, as it allows aliens to occupy niches that are not used by natives^79–81^. Here, we unveiled another role of novelty, which could decrease spill-over of endophytes between alien plant species, some of which are pathogenic. Consequently, alien species in our study suppressed each other less than they suppressed natives, and this could lead to invasional meltdown.

## Supporting information

all supplements

## Acknowledgements

We thank L. Arnold, S. Berg, O. Ficht, M. Fuchs, S. Gommel, E. Mamonova, V. Pasqualetto, C. Rabung, B. Rüter, B. Speißer, H. Vahlenkamp and E. Werner for practical assistance. ZZ was funded by the China Scholarship Council (201606100049) and supported by the International Max Planck Research School for Organismal Biology. YL was funded by the Chinese Academy of Sciences (Y9H1011001, Y9B7041001).

## Author contributions

ZZ conceived the idea. ZZ, YL and MvK designed the experiment. ZZ, YL and CB performed the experiment. ZZ analyzed the data and wrote the manuscript with input from all others.

