## Supplementary material for "Soil-microbes-mediated invasional meltdown in plants": all supplements

Supplement

### **Supplement S1 Information on plant species**

##### Table S1 The 13 species used in the experiment.

| **Species** | **Origin** | **Native origin** | **Family** | **Abbreviation** | **Grid**  **cells^b^** | **Sowing date soil-cond. phase** | **Transplanting date soil-conditioning phase** | **Sowing date test phase** |
| --- | --- | --- | --- | --- | --- | --- | --- | --- |
| *Epilobium ciliatum* | alien | North America | Onagraceae | - | 2448 | - | - | 9 Oct 2018 |
| *Lolium multiflorum* | alien^a^ | Europe | Poaceae | LM | 2639 | 27 June 2018 | 9 July 2018 | 18 Oct 2018 |
| *Onobrychis viciifolia* | alien^a^ | Europe | Leguminosae | OV | 1544 | 27 June 2018 | 9 July 2018 | 16 Oct 2018 |
| *Salvia verticillata* | alien^a^ | Europe | Lamiaceae | SV | 977 | 25 June 2018 | 25 July 2018 /  7 August 2018 | - |
| *Senecio inaequidens* | alien | Southern Africa | Compositae | - | 1322 | - | - | 16 Oct 2018 |
| *Solidago canadensis* | alien | North America | Compositae | - | 2660 | - | - | 9 Oct 2018 |
| *Solidago gigantea* | alien | North America | Compositae | SG | 2438 | 2 July 2018 | 30 July 2018/  12 August 2018 | - |
| *Cynosurus cristatus* | native | - | Poaceae | CC | 2710 | 27 June 2018 | 9 July 2018 | - |
| *Dactylis glomerata* | native | - | Poaceae | DG | 2990 | 27 June 2018 | 9 July 2018 | 16 Oct 2018 |
| *Leontodon autumnalis* | native | - | Compositae | LA | 2964 | 25 June 2018 | 9 July 2018 | 16 Oct 2018 |
| *Lotus corniculatus* | native | - | Leguminosae | LC | 2944 | 18 June 2018 | 9 July 2018 | 16 Oct 2018 |
| *Plantago media* | native | - | Plantaginaceae | PM | 2327 | 25 June 2018 | 11 July 2018 | 16 Oct 2018 |
| *Salvia pratensis* | native | - | Lamiaceae | SP | 1694 | 25 June 2018 | 9 July 2018 | 9 Oct 2018 |

Abbreviations are only for soil-conditioning species, and match the abbreviations in Fig. 4

^a^ intra-continental alien species (species that are alien in Germany and native in part of Europe)

^b^ the number of ~130-km^2^ grid cells occupied in Germany out of a maximum of 3000 grid cells (FloraWeb; <http://www.floraweb.de>).

### Supplement S2 Details on the statistical analyses

#### Analyses of plant performance

Model.plant.1

- Random effects: family and species identity of the test (focal and competitor plant) and soil-conditioning plants
- Covariate: transplanting date (to account for the fact that soil-conditioning plants were transplanted at different times)
- Variance structure: soil-conditioning species, test species and soil-conditioning treatments were allowed to vary their residuals by using the *varComb* and *varIdent* function^48^ (to improve homoscedasticity of residuals).
- Data transformation: aboveground biomass was square-root transformed (to improve normality of the residuals)

Model.plant.2

- Random effects: family and species identity of the test and soil-conditioning plants
- Covariate: transplanting date
- Variance structure: soil-conditioning species, test species and soil-conditioning treatments were allowed to vary their residuals
- Data transformation: aboveground biomass was square-root transformed (to improve normality of the residuals)

#### Analyses of the soil microbial community

Model.soil.1

- Random effects: family and species identity of the soil-conditioning plants (added as additional explanatory variables)
- Covariate: transplanting date (added as additional explanatory variables)
- Variance structure: not applicable (as this is PERMANOVA)
- Data transformation: no

Model.soil.2

- Random effects: family and species identity of soil-conditioning plants
- Covariate: transplanting date
- Variance structure: soil-conditioning species were allowed to have different variances (only for the models testing relative abundance)
- Data transformation: relative abundance was logit-transformed.

Model.soil.3

- Random effects: family and species identity of soil-conditioning species
- Covariate: no
- Variance structure: soil-conditioning species were allowed to have different variances
- Data transformation: Bray-Curtis dissimilarities were logit-transformed.

Model.link.1

- Random effects: family and species identity of soil-conditioning and test plants
- Covariate: none
- Variance structure: soil-conditioning and test species were allowed to have different variances (only for the models testing relative abundance)
- Data transformation: relative abundance was logit-transformed.

Model.link.2

- Random effects: family and species identity of soil-conditioning and test plants
- Covariate: none
- Variance structure: soil-conditioning and test species were allowed to have different variances
- Data transformation: Bray-Curtis dissimilarities were logit-transformed.

### Supplement S3 Results for plants.

##### Table S2 Effects of soil treatments, competition treatments, origin of test species and their interactions on aboveground biomass of plants after removing data where competitor test species were the same as the soil-conditioning species.

Significant effects (P < 0.05) are in bold and marked with asterisks, and marginally significant effects (0.05 ≤ P < 0.1) are in italics and marked with a dagger symbol.

|  | **χ^2^** | **P** |
| --- | --- | --- |
| Transplanting date | 12.693 | **<0.001*** |
| Soil_Non-conditioned/Conditioned_ | 10.624 | **0.001*** |
| Soil_Home/Away_ | 4.079 | **0.043*** |
| Soil_Alien/Native_ | 0.080 | 0.778 |
| Origin (O) | 0.086 | 0.770 |
| Comp_Yes/No_ | 3.681 | *0.055†* |
| Comp_Intra/Inter_ | 0.479 | 0.489 |
| O : Soil_Conditioned/Non-conditioned_ | 1.204 | 0.273 |
| O : Soil_Home/Away_ | 1.979 | 0.160 |
| O : Soil_Alien/Native_ | 6.338 | **0.012*** |
| Soil_Non-conditioned/Conditioned_: Comp_Yes/No_ | 0.906 | 0.341 |
| Soil_Non-conditioned/Conditioned_: Comp_Intra/Inter_ | 0.256 | 0.613 |
| Soil_Home/Away_ : Comp_Yes/No_ | 0.107 | 0.744 |
| Soil_Home/Away_ : Comp_Intra/Inter_ | 2.003 | 0.157 |
| Soil_Alien/Native_ : Comp_Yes/No_ | 5.497 | **0.019*** |
| Soil_Alien/Native_ : Comp_Intra/Inter_ | 0.127 | 0.721 |
| O:Comp_Yes/No_ | 0.275 | 0.600 |
| O:Comp_Intra/Inter_ | 0.395 | 0.529 |
| O : Soil_Non-conditioned/Conditioned_: Comp_Yes/No_ | 0.447 | 0.504 |
| O : Soil_Non-conditioned/Conditioned_: Comp_Intra/Inter_ | 0.001 | 0.979 |
| O : Soil_Home/Away_ : Comp_Yes/No_ | 1.751 | 0.186 |
| O : Soil_Home/Away_ : Comp_Intra/Inter_ | 0.260 | 0.610 |
| O : Soil_Alien/Native_ : Comp_Yes/No_ | 0.079 | 0.779 |
| O : Soil_Alien/Native_ : Comp_Intra/Inter_ | 0.080 | 0.777 |
| **Random effects** | SD |  |
| Family (focal test) | 0.164 |  |
| Species (focal test) | 0.200 |  |
| Family (competitor test) | 0.065 |  |
| Species (competitor test) | 0.076 |  |
| Family (soil) | 0.035 |  |
| Species (soil) | 0.033 |  |
| Residual | 0.179 |  |

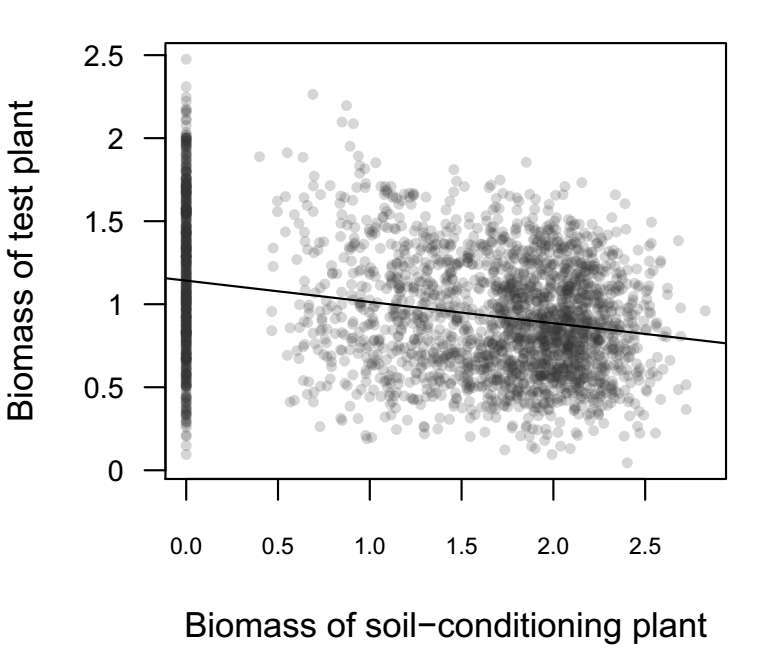

##### **Figure S1** Effects of aboveground biomass of soil-conditioning plant on aboveground biomass of test plant. Aboveground biomass of the empty control pots in the conditioning phase was set to 0.

##### Table S3 Effects of soil treatments, competition treatments, origin of test species and their interactions on aboveground biomass of plants (with square-root-transformed aboveground biomass of soil-conditioning plants as covariate).

|  | **χ^2^** | **P** |
| --- | --- | --- |
| Transplanting date | 0.917 | 0.338 |
| Sqrt(biomass_soil_) | 46.489 | **0.000*** |
| Soil_Non-conditioned/Conditioned_ | 4.758 | **0.029*** |
| Soil_Home/Away_ | 4.627 | **0.031*** |
| Soil_Alien/Native_ | 0.102 | 0.749 |
| Origin (O) | 0.077 | 0.781 |
| Comp_Yes/No_ | 3.722 | *0.054†* |
| Comp_Intra/Inter_ | 0.948 | 0.33 |
| O : Soil_Non-conditioned/Conditioned_ | 1.72 | 0.19 |
| O : Soil_Home/Away_ | 2.781 | *0.095†* |
| O : Soil_Alien/Native_ | 3.918 | **0.048*** |
| Soil_Non-conditioned/Conditioned_: Comp_Yes/N__o_ | 0.68 | 0.409 |
| Soil_Non-conditioned/Conditioned_: Comp_Intra/Inter_ | 0.098 | 0.754 |
| Soil_Home/Away_ : Comp_Yes/No_ | 0.248 | 0.619 |
| Soil_Home/Away_ : Comp_Intra/Inter_ | 1.551 | 0.213 |
| Soil_Alien/Native_ : Comp_Yes/No_ | 6.187 | **0.013*** |
| Soil_Alien/Native_ : Comp_Intra/Inter_ | 0.588 | 0.443 |
| O:Comp_Yes/No_ | 0.353 | 0.552 |
| O:Comp_Intra/Inter_ | 0.397 | 0.528 |
| O : Soil_Non-conditioned/Conditioned_: Comp_Yes/No_ | 0.485 | 0.486 |
| O : Soil_Non-conditioned/Conditioned_: Comp_Intra/Inter_ | 0.003 | 0.956 |
| O : Soil_Home/Away_ : Comp_Yes/No_ | 1.737 | 0.188 |
| O : Soil_Home/Away_ : Comp_Intra/Inter_ | 0.107 | 0.744 |
| O : Soil_Alien/Native_ : Comp_Yes/No_ | 0.124 | 0.725 |
| O : Soil_Alien/Native_ : Comp_Intra/Inter_ | 0.062 | 0.803 |
| **Random effects** | SD |  |
| Family (focal test) | 0.164 |  |
| Species (focal test) | 0.198 |  |
| Family (competitor test) | 0.065 |  |
| Species (competitor test) | 0.077 |  |
| Family (soil) | 0.000 |  |
| Species (soil) | 0.013 |  |
| Residual | 0.164 |  |

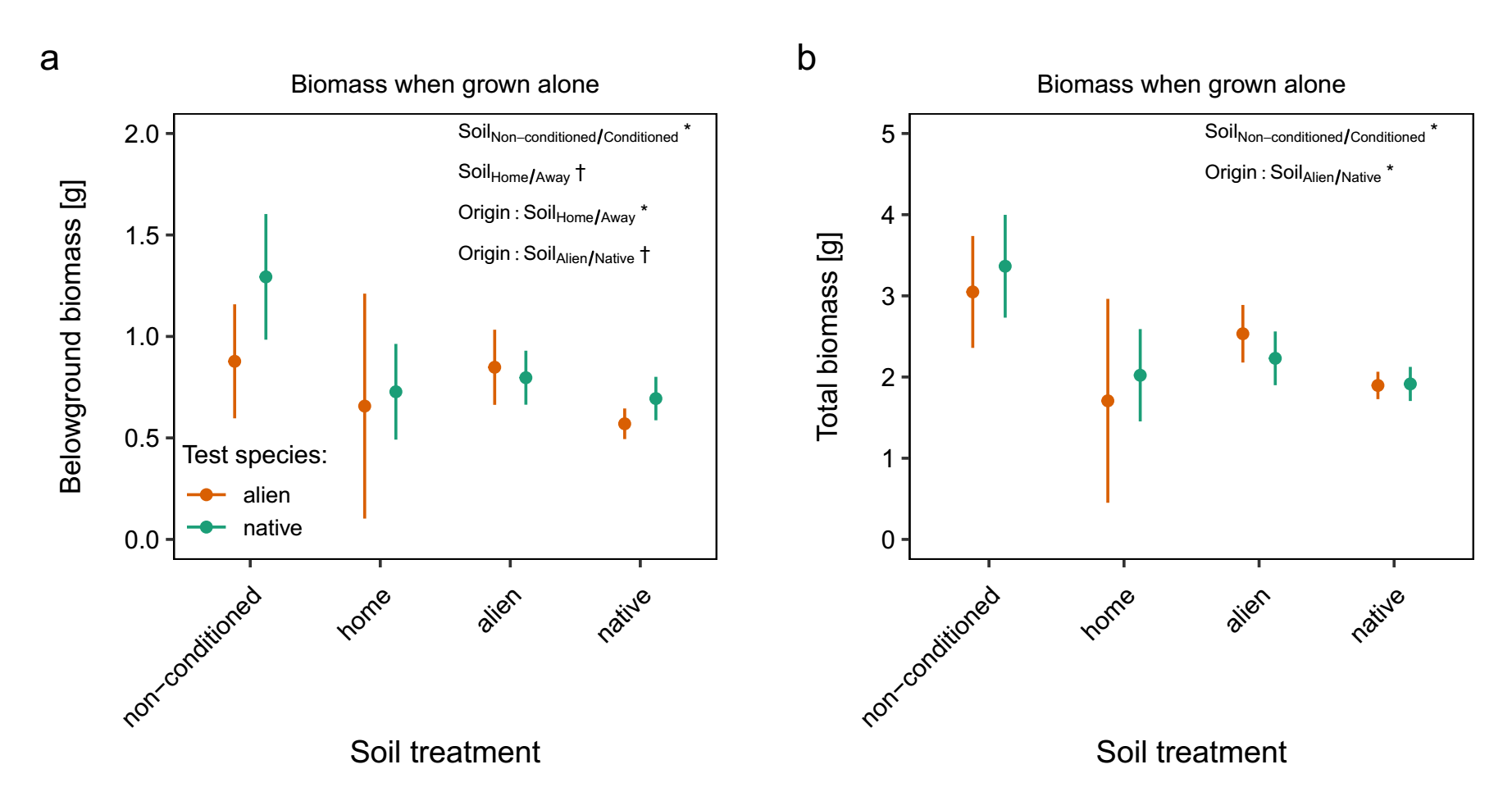

##### Figure S2 Effects of soil treatments on (a) belowground and (b) total biomass of alien (orange) and native (green) test species that were grown alone.

##### Table S4 Effects of soil treatments, competition treatments, origin of test species and their interactions on biomass of plants that were grown without competition.

Significant effects (P < 0.05) are in bold and marked with asterisks, and marginally significant effects (0.05 < P < 0.1) are in italics and marked with a dagger symbol.

|  | Aboveground biomass | |  | Belowground biomass | |  | Total biomass | |
| --- | --- | --- | --- | --- | --- | --- | --- | --- |
|  | χ^2^ | P |  | χ^2^ | P |  | χ^2^ | P |
| Transplanting date | 13.379 | **<0.001*** |  | 7.596 | **0.006*** |  | 13.265 | **<0.001*** |
| Soil_Non-conditioned/Conditioned_ | 7.618 | **0.006*** |  | 11.756 | **0.001*** |  | 8.430 | **0.004*** |
| Soil_Home/Away_ | 0.176 | 0.675 |  | 2.929 | *0.087†* |  | 1.834 | 0.176 |
| Soil_Alien/Native_ | 0.180 | 0.672 |  | 0.874 | 0.350 |  | 0.189 | 0.663 |
| Origin (O) | 0.025 | 0.875 |  | 1.995 | 0.158 |  | 0.229 | 0.632 |
| O : Soil_Non-conditioned/Conditioned_ | 0.076 | 0.783 |  | 5.439 | **0.020*** |  | 1.263 | 0.261 |
| O : Soil_Home/Away_ | 0.021 | 0.884 |  | 0.150 | 0.699 |  | 0.130 | 0.718 |
| O : Soil_Alien/Native_ | 2.902 | *0.088†* |  | 3.232 | *0.072†* |  | 4.563 | **0.033*** |
| **Random effects** | SD |  |  | SD |  |  | SD |  |
| Family (test) | 0.191 |  |  | 0.240 |  |  | 0.314 |  |
| Species (test) | 0.223 |  |  | 0.098 |  |  | 0.193 |  |
| Family (soil) | 0.056 |  |  | 0.000 |  |  | 0.053 |  |
| Species (soil) | 0.055 |  |  | 0.019 |  |  | 0.060 |  |
| Residual | 0.293 |  |  | 0.251 |  |  | 0.327 |  |

### Supplement S4 Effects of soil-conditioning plants on soil microbial communities

Supplement S4 is used to test whether and how soil-conditioning plants affected soil microbial communities (i.e. α in Fig. 1)

In addition to the main experiment, we had another 49 pots (five replicates for pots without plants and three to five replicates for each species) in which the field-soil inoculum was autoclaved before soil-conditioning. We used these pots to test the effect of the field soil, which provided the background soil microbial inoculum for the conditioning phase. Here, we present the results of both conditioned soils with live inoculum and conditioned soils with sterilized inoculum. For the relationship between soil communities and soil-legacy effects (supplement S4), we only used data from conditioned soils that had received live inoculum at the start. This is because test plants were grown on these soil and not on those with sterilized inoculum.

We detected a total of 45,288 bacterial phylotypes and 2,421 fungal phylotypes across all samples. Fungal functional groups (e.g. AMF, plant pathogens and endophytes) could be assigned to 889 phylotypes, representing 27.2 % of the ITS sequence reads. Of these phylotypes, 50 were identified as AMF, 165 as endophytes and 250 as plant pathogens, representing 0.1 %, 10.3%, and 15.1% of the ITS sequence reads, respectively.

#### S4.1 Diversity and composition of soil bacterial and fungal communities.

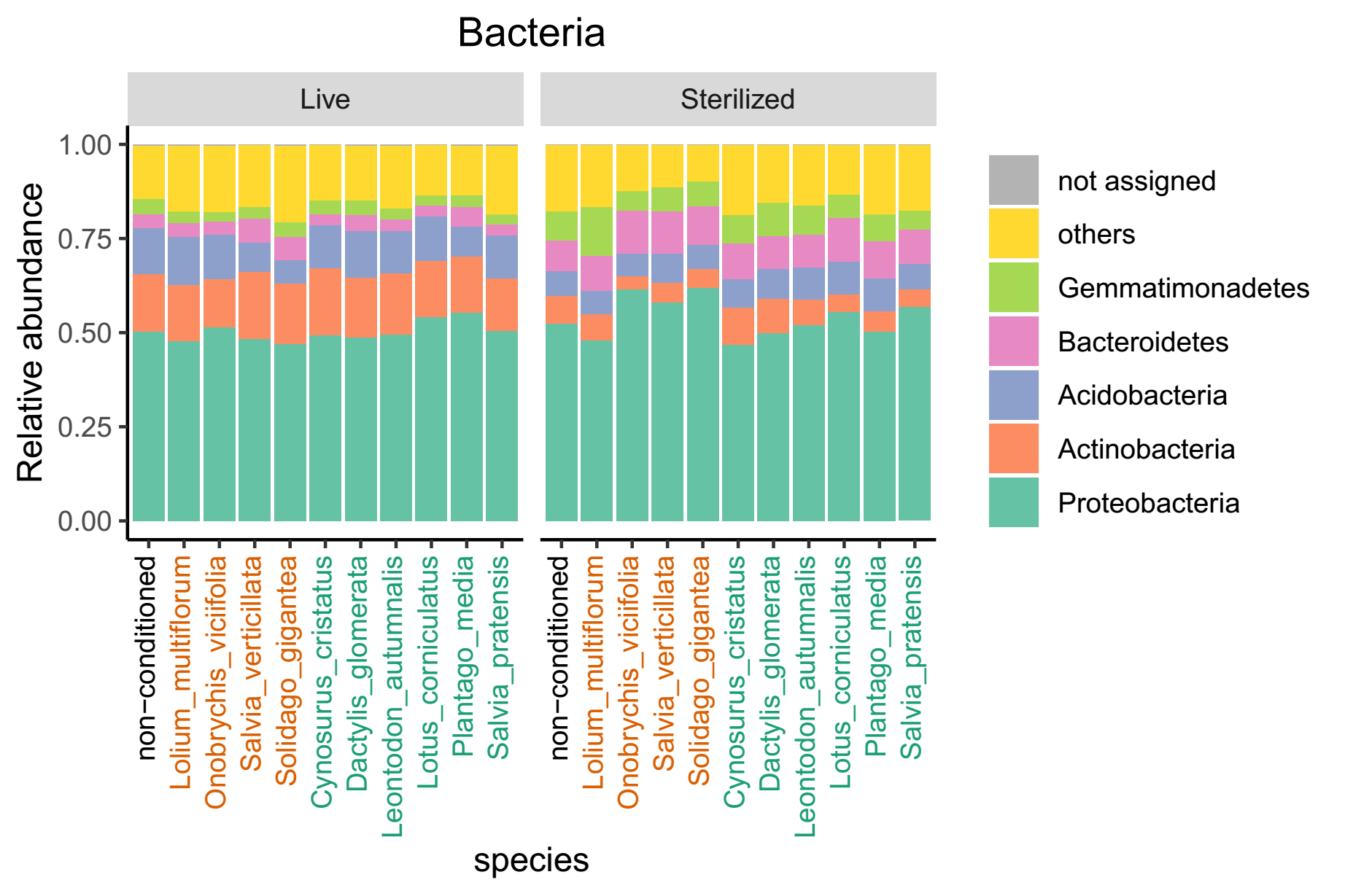

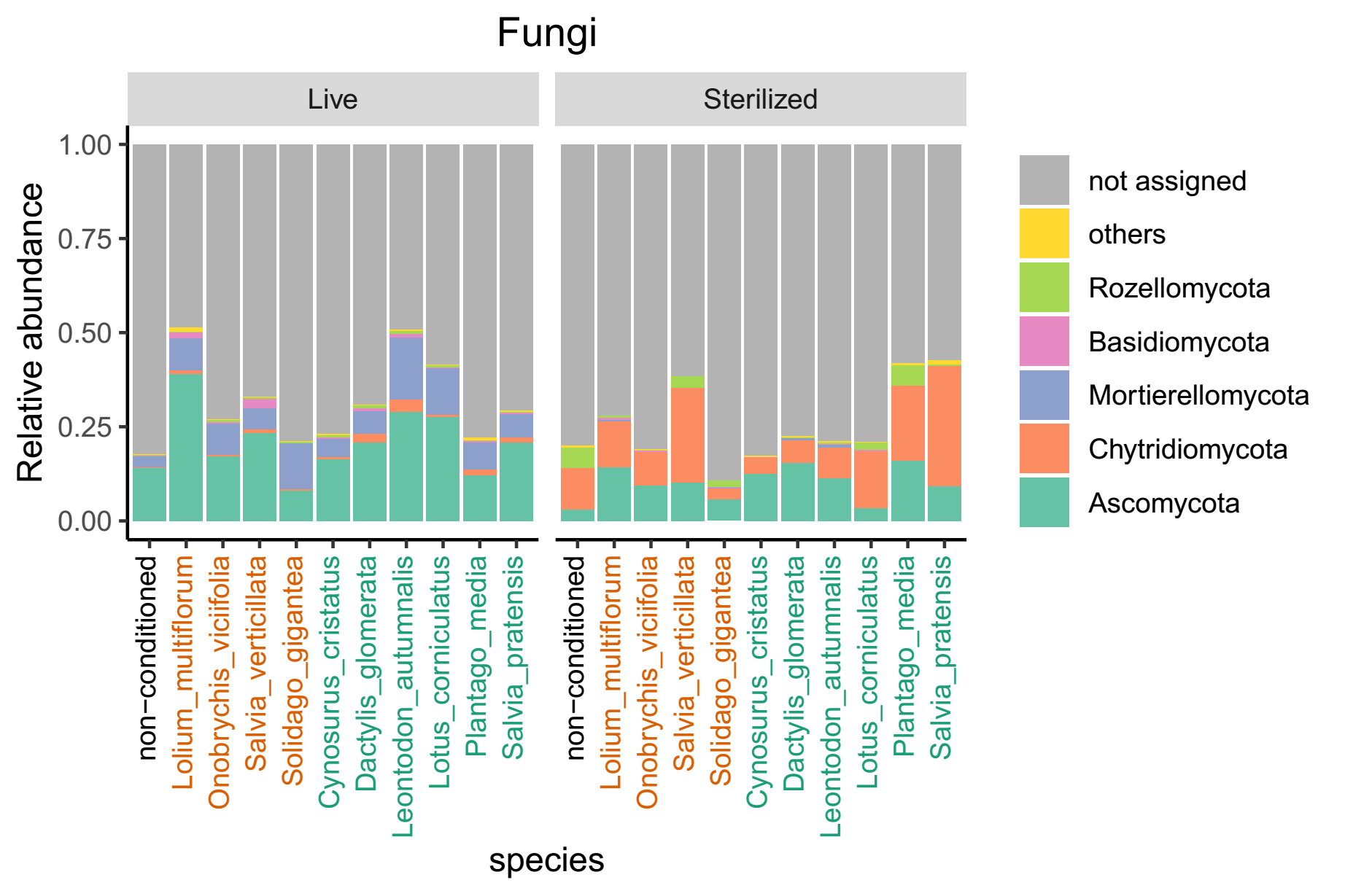

##### Figure S3 Relative abundances of major taxonomic groups of bacteria and fungi.

Alien and native plant species are in orange and green, respectively. Sterilized: field soil inoculum was autoclaved before soil-conditioning. Live: field soil was not treated before soil conditioning.
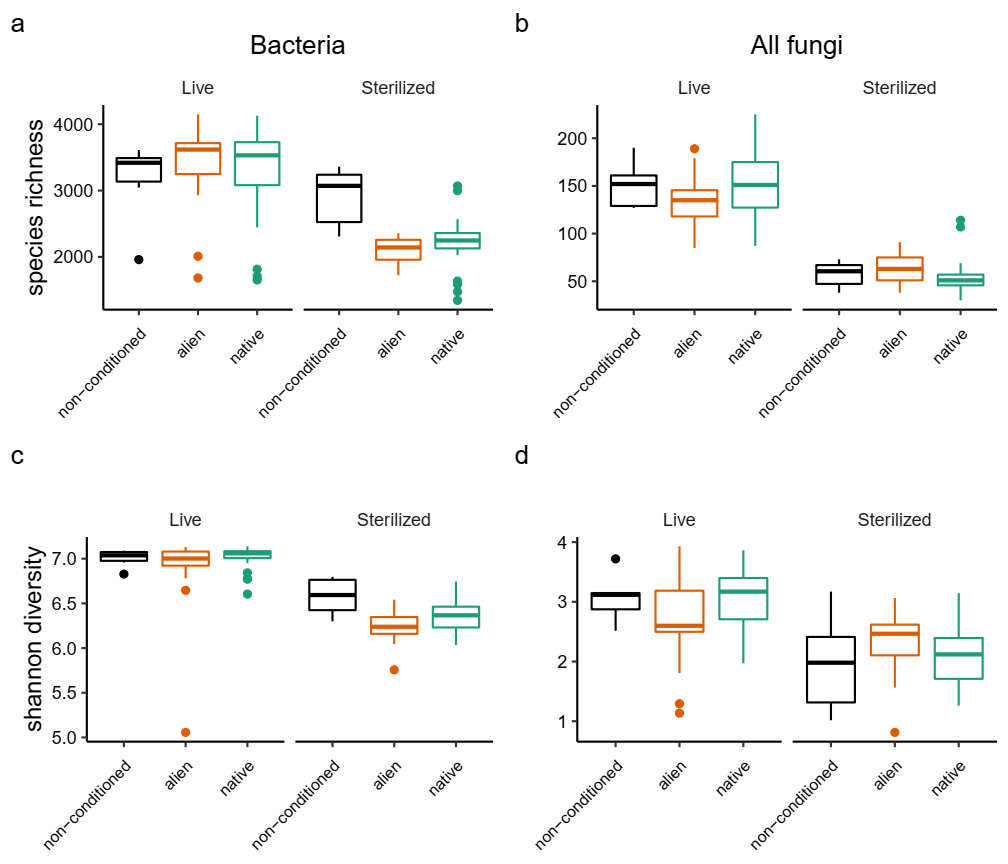

##### Figure S4 Effects of soil treatments on diversity of soil bacteria and fungi.

##### Table S5 Effects of soil treatments on diversity of soil bacteria and fungi.

Significant effects (P < 0.05) are in bold and marked with asterisks, and marginally significant effects (0.05 < P < 0.1) are in italics and marked with a dagger symbol.

Species richness

|  | Bacteria | | | | |  |  | All fungi | | |  |
| --- | --- | --- | --- | --- | --- | --- | --- | --- | --- | --- | --- |
|  | Live | |  | Sterilized | |  | Lived | |  | Sterilized | |
|  | χ^2^ | P |  | χ^2^ | P |  | χ^2^ | P |  | χ^2^ | P |
| Transplanting date | 0.298 | 0.585 | | 0.174 | 0.676 |  | 7.911 | **0.005*** |  | 0.000 | 0.988 |
| Soil_Conditioned/Non-conditioned_ | 0.647 | 0.421 |  | 13.548 | **0.000*** |  | 0.058 | 0.809 |  | 0.392 | 0.531 |
| Soil_Alien/Native_ | 0.180 | 0.671 |  | 0.795 | 0.373 |  | 0.386 | 0.535 |  | 4.368 | **0.037*** |
| **Random effects** | SD |  |  | SD |  |  | SD |  |  | SD |  |
| Family | 0.045 |  |  | 0.014 |  |  | 0.001 |  |  | 7.455 |  |
| Species | 0.057 |  |  | 0.016 |  |  | 0.001 |  |  | 0.001 |  |
| Residual | 601.913 |  |  | 336.203 |  |  | 28.912 |  |  | 15.038 |  |

Shannon diversity

|  | Bacteria | | | | |  |  | All fungi | | |  |
| --- | --- | --- | --- | --- | --- | --- | --- | --- | --- | --- | --- |
|  | Live | |  | Sterilized | |  | Lived | |  | Sterilized | |
|  | χ^2^ | P |  | χ^2^ | P |  | χ^2^ | P |  | χ^2^ | P |
| Transplanting date | 5.329 | **0.021*** | | 2.561 | 0.109 |  | 2.789 | *0.095†* |  | 0.588 | 0.443 |
| Soil_Conditioned/Non-conditioned_ | 0.001 | 0.982 |  | 8.389 | **0.004*** |  | 0.260 | 0.610 |  | 0.841 | 0.359 |
| Soil_Alien/Native_ | 0.211 | 0.646 |  | 2.215 | 0.137 |  | 1.375 | 0.241 |  | 0.983 | 0.322 |
| **Random effects** | SD |  |  | SD |  |  | SD |  |  | SD |  |
| Family | 0 |  |  | 0 |  |  | 0 |  |  | 0 |  |
| Species | 0 |  |  | 0 |  |  | 0 |  |  | 0 |  |
| Residual | 0.255 |  |  | 0.172 |  |  | 0.588 |  |  | 0.542 |  |

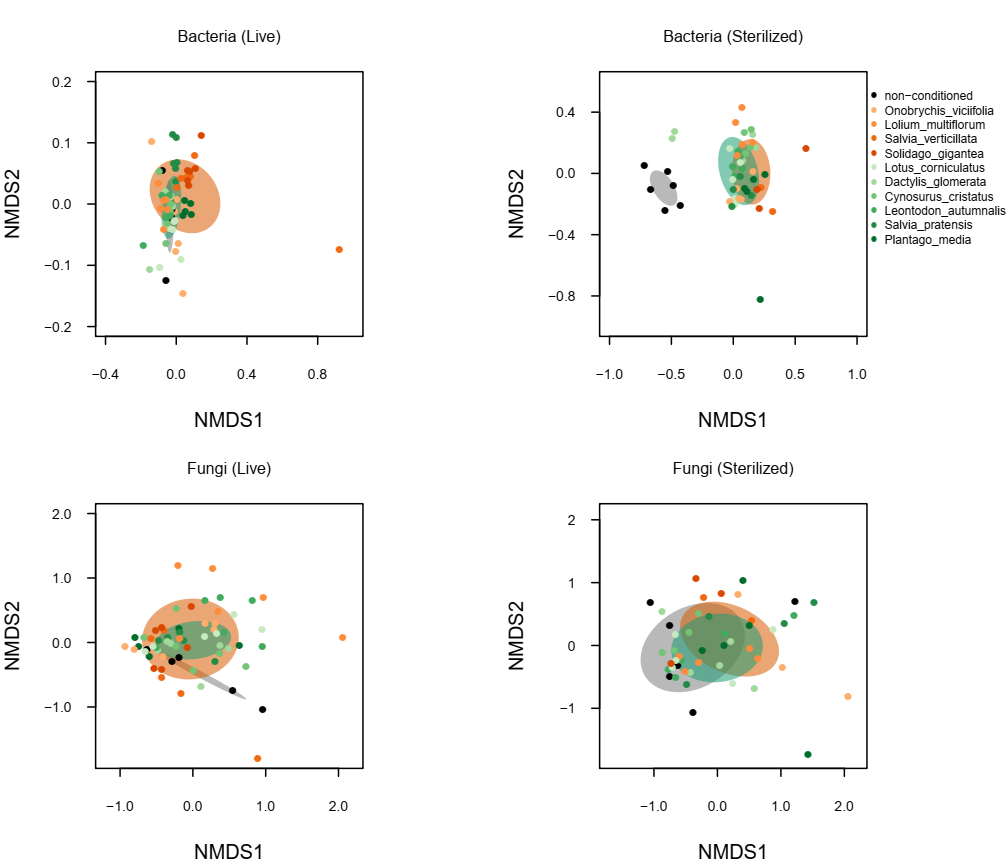

##### Figure S5 Soil community compositions of bacteria and fungi.

We used nonmetric multidimensional scaling (NMDS) to visualize differences in the soil microbial communities of the plant species. Data points represent soil samples. Ellipses represent means ± 1 SDs for soil conditioned by aliens (orange) or natives (green), or not conditioned by plants (grey).

##### Table S6 Effects of soil treatments on composition of soil bacterial and fungi communities.

Significant effects (P < 0.05) are in bold and marked with asterisks, and marginally significant effects (0.05 ≤ P < 0.1) are in italics and marked with a dagger symbol.

|  | Live | | | |  | Sterilized | | | |
| --- | --- | --- | --- | --- | --- | --- | --- | --- | --- |
|  | Df | F | R^2^ | P |  | Df | F | R^2^ | P |
| Transplanting date | 1 | 1.226 | 0.016 | 0.113 |  | - | - | - | - |
| Soil_Conditioned/Non-conditioned_ | 1 | 1.711 | 0.022 | **0.006*** |  | 1 | 8.087 | 0.124 | **0.001*** |
| Soil_Alien/Native_ | 1 | 2.442 | 0.032 | **0.001*** |  | 1 | 2.298 | 0.035 | **0.003*** |
| Family | 4 | 2.790 | 0.146 | **0.001*** |  | 4 | 2.705 | 0.165 | **0.001*** |
| Species | 4 | 1.684 | 0.088 | **0.001*** |  | 4 | 1.553 | 0.095 | **0.002*** |
| Residuals | 53 |  | 0.695 |  |  | 38 |  | 0.581 |  |
| Total | 64 |  | 1.000 |  |  | 48 |  | 1.000 |  |

Bacteria

Fungi

|  | Live | | | |  | Sterilized | | | |
| --- | --- | --- | --- | --- | --- | --- | --- | --- | --- |
|  | Df | F | R^2^ | P |  | Df | F | R^2^ | P |
| Transplanting date | 1 | 1.148 | 0.017 | 0.275 |  | - | - | - | - |
| Soil_Conditioned/Non-conditioned_ | 1 | 1.493 | 0.023 | *0.076†* |  | 1 | 1.198 | 0.024 | 0.263 |
| Soil_Alien/Native_ | 1 | 1.826 | 0.028 | **0.025*** |  | 1 | 1.498 | 0.031 | 0.109 |
| Family | 4 | 1.361 | 0.082 | **0.044*** |  | 4 | 1.352 | 0.111 | *0.074†* |
| Species | 4 | 1.517 | 0.092 | **0.006*** |  | 4 | 1.198 | 0.098 | 0.164 |
| Residuals | 50 |  | 0.758 |  |  | 36 |  | 0.736 |  |
| Total | 61 |  | 1.000 |  |  | 46 |  | 1.000 |  |

For samples whose field soil inoculum was sterilized, effects of transplanting date cannot be tested, because we transplanted *S. verticillata* and *S. gigantean* only once there. Consequently, collinearity between transplanting date and species prevented us from including both factors in the same models.

#### S4.2 Abundance, diversity and composition of fungal pathogen communities

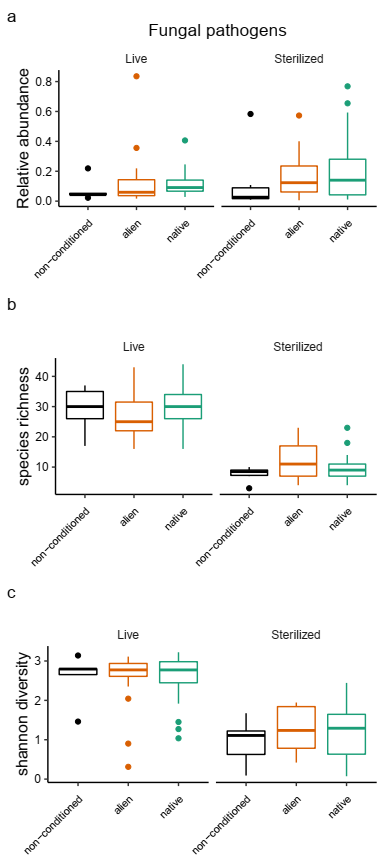

##### Figure S6 Effects of soil treatments on relative abundance and diversity of fungal pathogens.

##### Table S7 Effects of soil treatments on relative abundance and diversity of soil fungal pathogens.

Significant effects (P < 0.05) are in bold and marked with asterisks, and marginally significant effects (0.05 < P < 0.1) are in italics and marked with a dagger symbol.

Relative abundance

|  | Pathogens | | | | |
| --- | --- | --- | --- | --- | --- |
|  | Live | |  | Sterilized | |
|  | χ^2^ | P |  | χ^2^ | P |
| Transplanting date | 11.932 | **0.001*** | | 2.800 | *0.094†* |
| Soil_Conditioned/Non-conditioned_ | 1.325 | 0.250 |  | 2.537 | 0.111 |
| Soil_Alien/Native_ | 0.002 | 0.967 |  | 1.379 | 0.240 |
| **Random effects** | SD |  |  | SD |  |
| Family | 0.000 |  |  | 0.000 |  |
| Species | 0.000 |  |  | 0.000 |  |
| Residual | 1.043 |  |  | 0.843 |  |

Species richness

|  | Pathogens | | | | |
| --- | --- | --- | --- | --- | --- |
|  | Live | |  | Sterilized | |
|  | χ^2^ | P |  | χ^2^ | P |
| Transplanting date | 8.487 | **0.004*** | | 2.601 | 0.107 |
| Soil_Conditioned/Non-conditioned_ | 0.144 | 0.704 |  | 4.250 | **0.039*** |
| Soil_Alien/Native_ | 0.008 | 0.930 |  | 6.707 | **0.010*** |
| **Random effects** | SD |  |  | SD |  |
| Family | 0.000 |  |  | 0.000 |  |
| Species | 1.585 |  |  | 0.938 |  |
| Residual | 5.975 |  |  | 3.807 |  |

Shannon diversity

|  | Pathogens | | | | |
| --- | --- | --- | --- | --- | --- |
|  | Live | |  | Sterilized | |
|  | χ^2^ | P |  | χ^2^ | P |
| Transplanting date | 0.364 | 0.546 |  | 2.911 | *0.088†* |
| Soil_Conditioned/Non-conditioned_ | 0.038 | 0.845 |  | 1.390 | 0.238 |
| Soil_Alien/Native_ | 0.002 | 0.966 |  | 2.340 | 0.126 |
| **Random effects** | SD |  |  | SD |  |
| Family | 0.000 |  |  | 0.151 |  |
| Species | 0.000 |  |  | 0.000 |  |
| Residual | 0.587 |  |  | 0.542 |  |

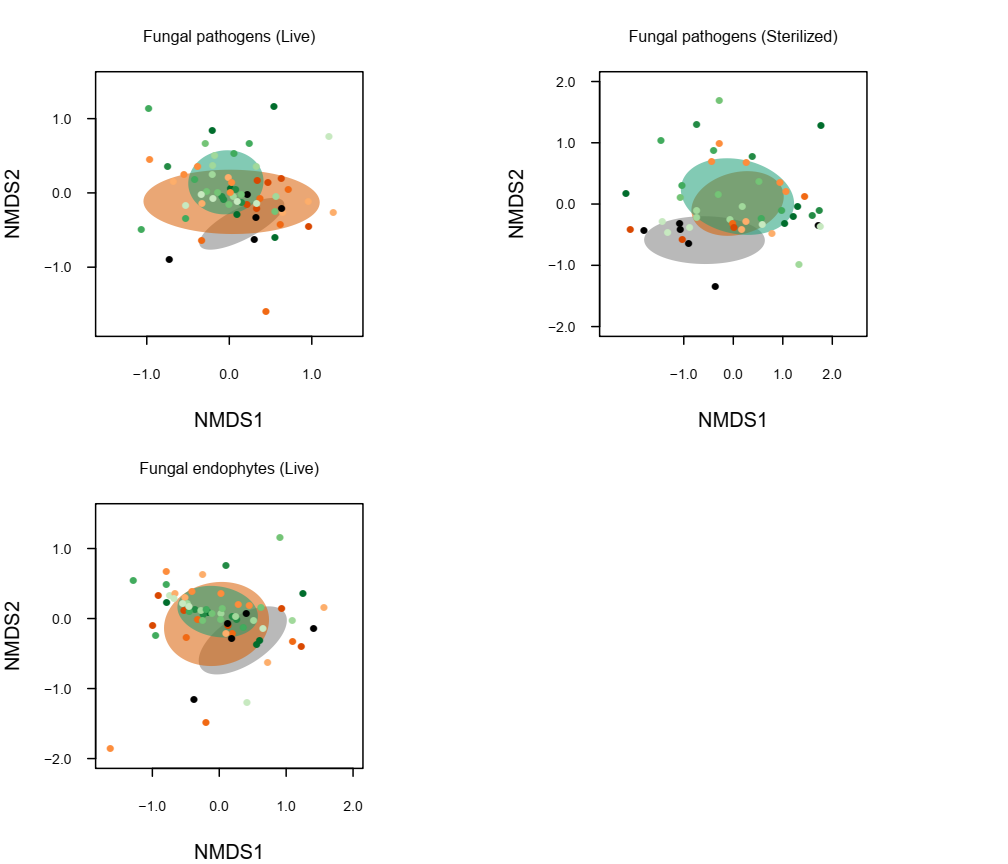

##### Figure S7 Differences in composition of fungal pathogen soil communities.

Data points represent soil samples. Ellipses represent means ± 1 SDs for soils conditioned by aliens (orange) or natives (green), or not conditioned by plants (grey). The different shades of the colors used for the point indicate different species. See Figure S3 for details.

##### Table S8 Effects of soil treatments on community composition of soil fungal pathogens.

Significant effects (P < 0.05) are in bold and marked with asterisks, and marginally significant effects (0.05 < P < 0.1) are in italics and marked with a dagger symbol.

|  | Live | | | |  | Sterilized | | | |
| --- | --- | --- | --- | --- | --- | --- | --- | --- | --- |
|  | Df | F | R^2^ | P |  | Df | F | R^2^ | P |
| Transplanting date | 1 | 0.851 | 0.013 | 0.668 |  | - | - | - | - |
| Soil_Conditioned/Non-conditioned_ | 1 | 1.387 | 0.021 | *0.097†* |  | 1 | 2.379 | 0.047 | **0.022*** |
| Soil_Alien/Native_ | 1 | 1.995 | 0.031 | **0.011*** |  | 1 | 0.497 | 0.010 | 0.910 |
| family | 4 | 1.228 | 0.076 | *0.084†* |  | 4 | 1.611 | 0.128 | **0.029*** |
| species | 4 | 1.475 | 0.091 | **0.009*** |  | 4 | 1.302 | 0.103 | 0.127 |
| Residuals | 50 |  | 0.769 |  |  | 36 |  | 0.713 |  |
| Total | 61 |  | 1.000 |  |  | 46 |  | 1.000 |  |

#### S4.3 Dissimilarity of soil communities

##### Table S9 Dissimilarities of soil communities within and between species.

Four contrasts were made to address the questions: 1) Are soil communities more similar (or different) when conditioned by the same plant species than by a different species? 2) When conditioned by the same species, are soil communities more similar when conditioned by alien species than by native species? 3) When conditioned by different species, are soil communities more similar between two aliens than between an alien and a native species? 4) When conditioned by different species, are soil communities more similar between an alien and a native species than between two natives. Significant effects (P < 0.05) are in bold and marked with asterisks, and marginally significant effects (0.05 < P < 0.1) are in italics and marked with a dagger symbol.

|  | Bacteria | |  | All fungi | |  | Fungal pathogens | |  | Fungal endophytes | |
| --- | --- | --- | --- | --- | --- | --- | --- | --- | --- | --- | --- |
|  | χ^2^ | P |  | χ^2^ | P |  | χ^2^ | P |  | χ^2^ | P |
| Within-species vs between species | 4.307 | **0.038*** |  | 1.038 | 0.308 |  | 2.004 | 0.157 |  | 0.150 | 0.698 |
| Within-species: alien vs native | 0.080 | 0.777 |  | 1.122 | 0.289 |  | 2.271 | 0.132 |  | 16.292 | **0.000*** |
| Alien-alien vs alien-native | 0.706 | 0.401 |  | 4.001 | **0.045*** |  | 1.358 | 0.244 |  | 12.108 | **0.001*** |
| Alien-native vs native-native | 2.687 | 0.101 |  | 3.123 | *0.077†* |  | 2.271 | 0.132 |  | 10.527 | **0.001*** |
| **Random effects** | SD |  |  | SD |  |  | SD |  |  | SD |  |
| species1 | 0.079 |  |  | 0.137 |  |  | 0.160 |  |  | 0.104 |  |
| family1 | 0.063 |  |  | 0.000 |  |  | 0.000 |  |  | 0.000 |  |
| species2 | 0.006 |  |  | 0.207 |  |  | 0.195 |  |  | 0.058 |  |
| family2 | 0.042 |  |  | 0.071 |  |  | 0.000 |  |  | 0.022 |  |
| Residual | 0.080 |  |  | 0.039 |  |  | 0.130 |  |  | 0.120 |  |

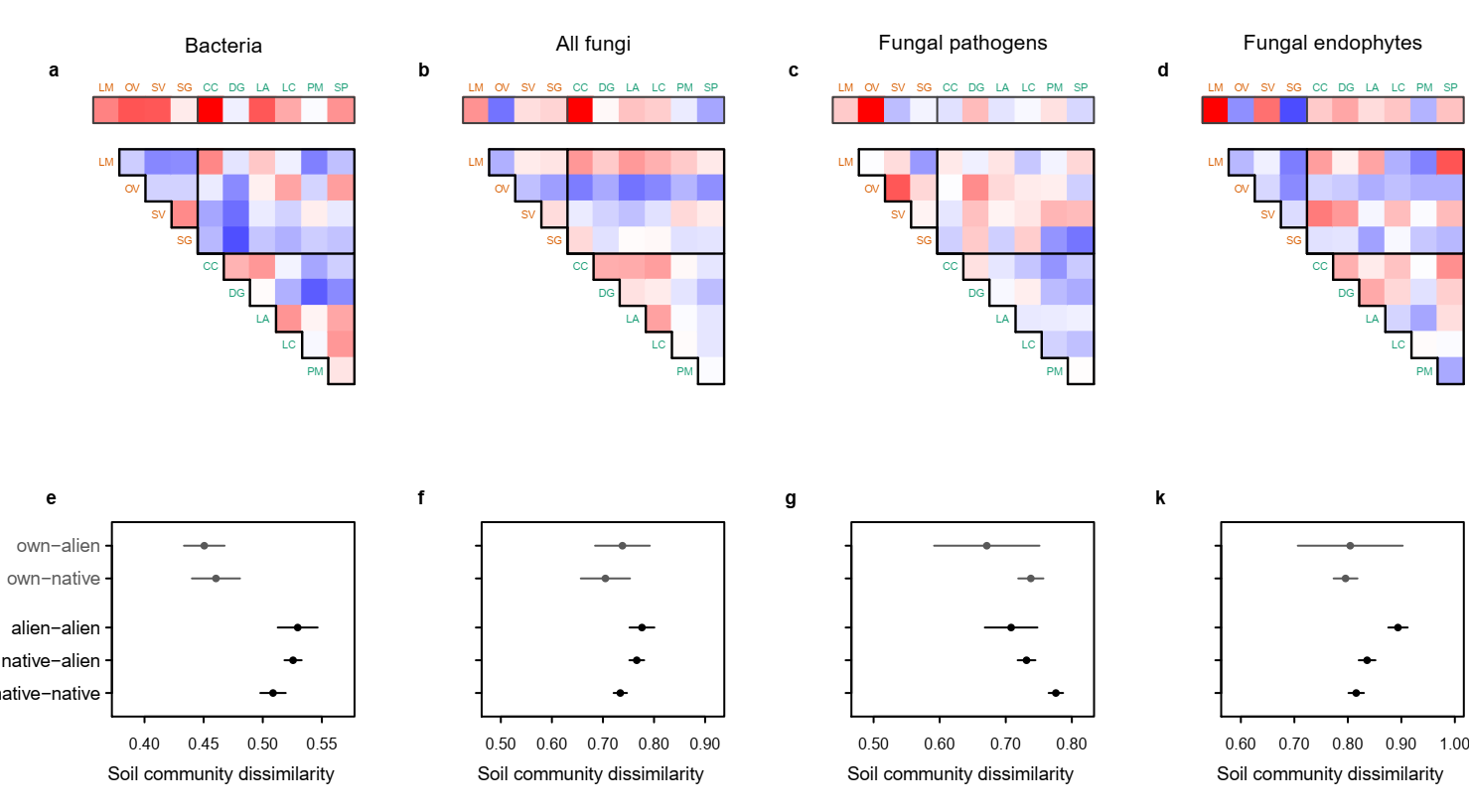

##### Figure S8 Dissimilarities of soil communities within and between species, when sterilized field soil was used as inoculum.

**a** & **e**, bacterial communities; **b** & **f**, fungal communities, **c** & g, fungal pathogen communities; **d** & **h**, fungal endophyte communities. The upper panel shows the heatmaps of community dissimilarities of all within-species (top horizontal bars) and between-species combinations (triangular matrices), which are divided into five categories (own-alien, own-native, alien-alien, native-alien, native-native) with black borders. Labels at the top and along the diagonal provide abbreviations of the names (see Table S1) of alien (orange) and native (green) species. The lower panel shows the mean values (±SEs) of each of the five categories. Own-alien: between plants of the same alien plant species; own-native: between plants of the same native species; alien-alien: between plants of different alien species; alien-native: between plants of alien and native species; native-native: between plants of different native species.

##### Table S10 Dissimilarities of soil communities within and between species, when sterilized field soil was used as inoculum.

Contrasts are the same as in table S8. Significant effects (P < 0.05) are in bold and marked with asterisks, and marginally significant effects (0.05 < P < 0.1) are in italics and marked with a dagger symbol.

|  | Bacteria | |  | All fungi | |  | Fungal pathogens | |  | Fungal endophytes | |
| --- | --- | --- | --- | --- | --- | --- | --- | --- | --- | --- | --- |
|  | χ^2^ | P |  | χ^2^ | P |  | χ^2^ | P |  | χ^2^ | P |
| Wthin-species vs between species | 16.693 | **<0.001*** |  | 0.069 | 0.793 |  | 0.589 | 0.443 |  | 0.508 | 0.476 |
| Within-species: alien vs native | 0.559 | 0.454 |  | 0.085 | 0.771 |  | 0.637 | 0.425 |  | 1.105 | 0.293 |
| Alien-alien vs alien-native | 0.002 | 0.962 |  | 0.090 | 0.764 |  | 0.563 | 0.453 |  | 2.159 | 0.142 |
| Alien-native vs native-native | 0.257 | 0.612 |  | 0.673 | 0.412 |  | 3.097 | *0.078†* |  | 0.543 | 0.461 |
| **Random effects** | SD |  |  | SD |  |  | SD |  |  | SD |  |
| species1 | 0.148 |  |  | 0.400 |  |  | 0.137 |  |  | 0.263 |  |
| family1 | 0.085 |  |  | 0.196 |  |  | 0.005 |  |  | 0.004 |  |
| species2 | 0.087 |  |  | 0.175 |  |  | 0.063 |  |  | 0.256 |  |
| family2 | 0.095 |  |  | 0.000 |  |  | 0.004 |  |  | 0.001 |  |
| Residual | 0.060 |  |  | 0.145 |  |  | 0.157 |  |  | 0.000 |  |

### Supplement S5 Effect of soil microbes on test plants when grown alone

Supplement S5 is used to test whether soil microbes explained the soil-legacy effect (β_alone_ in Fig.1, the effect of soil-conditioning species on test plant when grown alone)

#### S5.1 Effects of diversity of soil bacteria and fungi

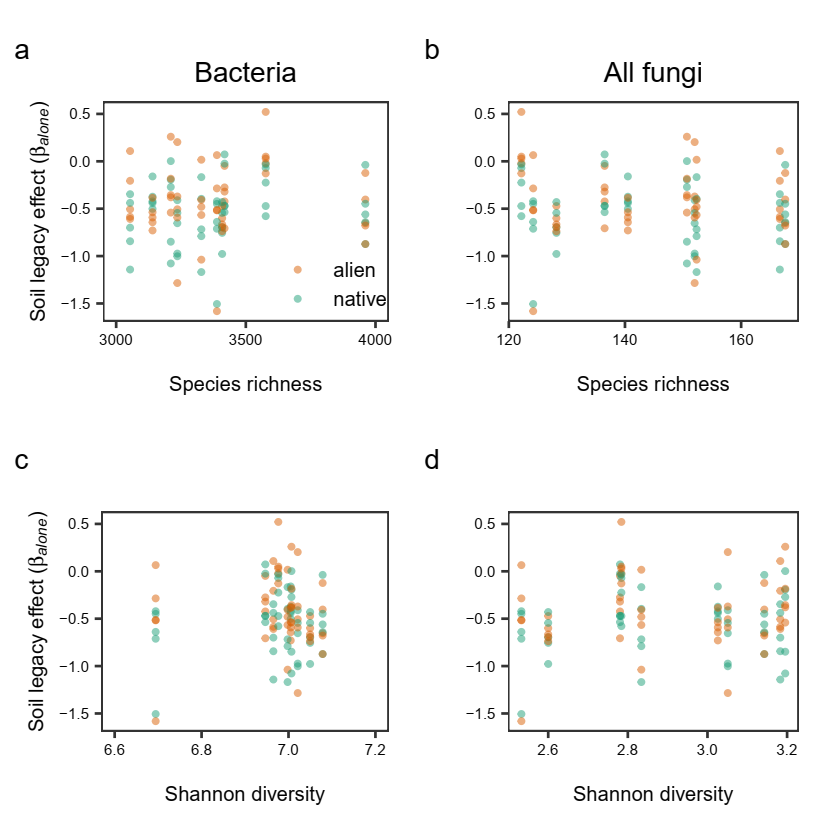

##### Figure S9 Effects of diversity (species richness or Shannon diversity) of bacteria (a, c) and fungi (b, d) on soil-legacy effects.

Green dots represent native test species, and orange dots represent alien test species. Negative values of soil-legacy effects indicate that plants grew worse on conditioned soil than on non-conditioned soil. No significant relationship was found.

##### Table S11 Effects of **diversity of bacteria and fungi on soil-legacy effects.**

Marginally significant effects (0.05 < P < 0.1) are in italics and marked with a dagger symbol.

Species richness

|  | Bacteria | |  | All fungi | |
| --- | --- | --- | --- | --- | --- |
|  | χ^2^ | P |  | χ^2^ | P |
| Species richness (SR) | 0.594 | 0.441 |  | 3.856 | *0.050†* |
| Origin (O) | 2.555 | 0.110 |  | 2.579 | 0.108 |
| O : SR | 0.509 | 0.475 |  | 0.501 | 0.479 |
| **Random effects** | SD |  |  | SD |  |
| Family (soil) | 0.156 |  |  | 0.175 |  |
| Species (soil) | 0.11 |  |  | 0.077 |  |
| Family (test) | 0.092 |  |  | 0.097 |  |
| Species (test) | 0.000 |  |  | 0.000 |  |
| Residual | 0.185 |  |  | 0.201 |  |

Shannon diversity

|  | Bacteria | |  | All fungi | |
| --- | --- | --- | --- | --- | --- |
|  | χ^2^ | P |  | χ^2^ | P |
| Shannon diversity (Sh) | 0.384 | 0.535 |  | 1.656 | 0.198 |
| Origin (O) | 2.576 | 0.108 |  | 2.761 | *0.097†* |
| O : Sh | 0.845 | 0.358 |  | 0.08 | 0.777 |
| **Random effects** | SD |  |  | SD |  |
| Family (soil) | 0.113 |  |  | 0.201 |  |
| Species (soil) | 0.135 |  |  | 0.060 |  |
| Family (test) | 0.097 |  |  | 0.097 |  |
| Species (test) | 0.000 |  |  | 0.000 |  |
| Residual | 0.231 |  |  | 0.199 |  |

#### S5.2 Effect of abundance and diversity of soil fungal pathogen

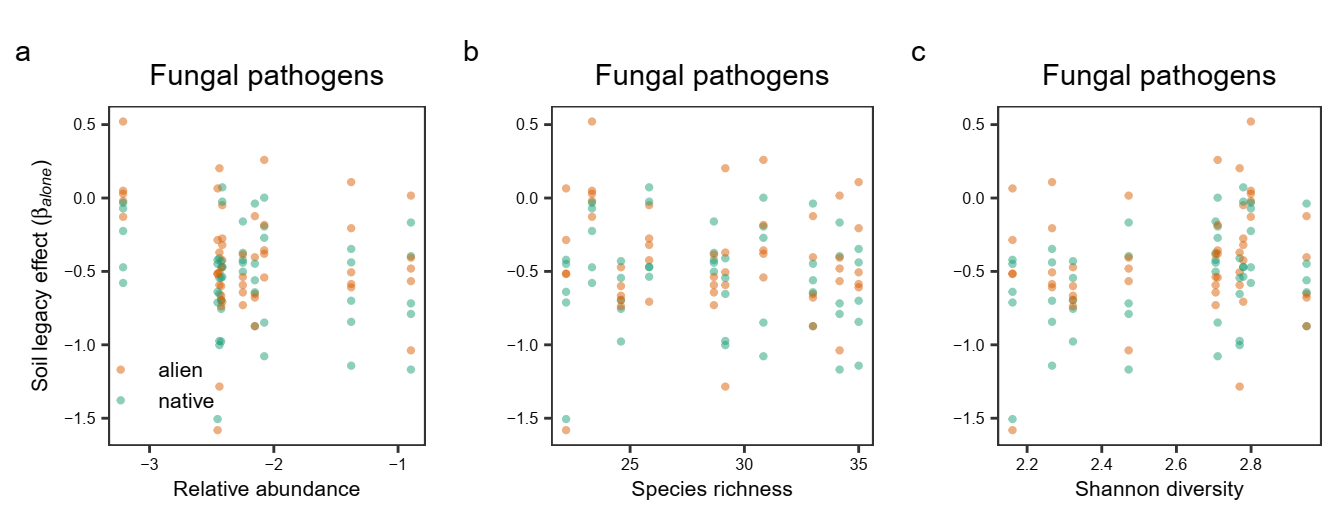

##### Figure S10 Effects of (a) relative abundance, (b) species richness and (c) Shannon diversity of fungal pathogens on soil-legacy effects.

Green dots represent native test species, and orange dots represent alien test species. Negative values of the soil-legacy effect indicate that plants grow worse on conditioned soil than on non-conditioned soil. No significant relationship was found.

##### Table S12 Effects of **relative abundance, species richness, or Shannon diversity of fungal pathogens on soil-legacy effects.**

Relative abundance

|  | Fungal pathogens | |
| --- | --- | --- |
|  | χ^2^ | P |
| Abundance | 2.448 | 0.118 |
| Origin (O) | 2.409 | 0.121 |
| O : abundance | 0.945 | 0.331 |
| **Random effects** | SD |  |
| Family (soil) | 0.135 |  |
| Species (soil) | 0.101 |  |
| Family (test) | 0.078 |  |
| Species (test) | 0.000 |  |
| Residual | 0.224 |  |

Species richness

|  | Fungal pathogens | |
| --- | --- | --- |
|  | χ^2^ | P |
| Species richness (SR) | 2.569 | 0.109 |
| Origin (O) | 2.555 | 0.110 |
| O : SR | 0.095 | 0.758 |
| **Random effects** | SD |  |
| Family (soil) | 0.161 |  |
| Species (soil) | 0.089 |  |
| Family (test) | 0.081 |  |
| Species (test) | 0.000 |  |
| Residual | 0.224 |  |

Shannon diversity

|  | Fungal pathogens | |
| --- | --- | --- |
|  | χ^2^ | P |
| Shannon diversity (Sh) | 1.296 | 0.255 |
| Origin (O) | 2.443 | 0.118 |
| O : Sh | 0.032 | 0.859 |
| **Random effects** | SD |  |
| Family (soil) | 0.091 |  |
| Species (soil) | 0.141 |  |
| Family (test) | 0.083 |  |
| Species (test) | 0.000 |  |
| Residual | 0.237 |  |

#### S5.3 Effect of soil-community dissimilarity on aboveground biomass

##### Table S13 Effects of soil community dissimilarity on soil-legacy effects.

Significant effects (P < 0.05) are in bold and marked with asterisks, and marginally significant effects (0.05 ≤ P < 0.1) are in italics and marked with a dagger symbol.

|  | Bacteria | |  | All fungi | |  | Fungal pathogens | |  | Fungal endophytes | |
| --- | --- | --- | --- | --- | --- | --- | --- | --- | --- | --- | --- |
|  | χ^2^ | P |  | χ^2^ | P |  | χ^2^ | P |  | χ^2^ | P |
| Logit(dissimilarity) | 2.777 | *0.096†* |  | 2.565 | 0.109 |  | -0.083 | 1.000 |  | 7.49 | **0.006*** |
| **Random effects** | SD |  |  | SD |  |  | SD |  |  | SD |  |
| Family(soil) | 0.186 |  |  | 0.184 |  |  | 0.160 |  |  | 0.193 |  |
| Species(soil) | 0.000 |  |  | 0.000 |  |  | 0.000 |  |  | 0.060 |  |
| Family(test) | 0.107 |  |  | 0.141 |  |  | 0.000 |  |  | 0.164 |  |
| Species(test) | 0.020 |  |  | 0.000 |  |  | 0.121 |  |  | 0.000 |  |
| Residual | 0.238 |  |  | 0.260 |  |  | 0.315 |  |  | 0.241 |  |

The analysis included intraspecific soil-legacy effects, i.e. the effect of one species on itself, for seven species. The evolutionary history of intraspecific soil-legacy effects may differ from that of interspecific soil-legacy effects. Therefore, we tried to add an intraspecific v.s. interspecific effect to account for this potential effect. However, the intraspecific v.s. interspecific effect was not significant and adding it did not affect the effect of dissimilarity. Therefore, we did not include it here.

### Supplement S6 Effects of soil microbes on the strength of competition between test plants

Supplement S6 describes the method of how we tested whether soil microbes explained the soil legacy effect (β_inter_ or β_intra_ in Fig.1, the effect of soil-conditioning species on strength of competition between focal test and competitor test species), and presents results.

**The method**

We calculated he soil-legacy effect on the strength of interspecific competition, $\beta_{inter, i,j,k}$, as:

$$\beta_{inter, i,j,k}={Strength}_{i,j,k}- {Strength}_{i,j,0} (i !=j)$$

$${Strength}_{i,j,k}= \ln{biomass}_{i,j, k}-\ln{biomass}_{i,0, k}$$

$${Strength}_{i,j,0}= \ln{biomass}_{i,j, 0}-\ln{biomass}_{i,0, 0}$$

Here, $\ln{biomass}_{i,j,k}$ is the mean aboveground biomass of focal test species *i* when grown with competitor test species *j* on **soil conditioned by species *k***. $\ln{biomass}_{i,0,k}$ is the mean aboveground biomass of focal test species *i* when grown alone on soil conditioned by species *k.* Negative values of ${Strength}_{i,j,k}$ indicate that on soil conditioned by species *k*, species *i* was negatively affected by species *j*.

Similarly, $\ln{biomass}_{i,j,0}$ is the mean aboveground biomass of focal test species *i* when grown with competitor test species *j* on **non-conditioned soil**. $\ln{biomass}_{i,0,0}$ is the mean aboveground biomass of focal test species *i* when grown alone on *non-conditioned soil.* Negative values of ${Strength}_{i,j,0}$ indicate that on *non-conditioned soil*, species *i* was negatively affected by species *j*.

Consequently, negative values of $\beta_{inter, i,j,k}$ indicate that the effects of competitor test species *j* on the focal test species *i* were more negative on soil conditioned by species *k* than on non-conditioned soil. In other words, the strength of interspecific competition was more negative on soil conditioned by species *k*.

Likewise, the soil-legacy effect on the strength of intraspecific competition, $\beta_{intra, i,i,k}$, was calculated as:

$$\beta_{intra, i,i,k}={Strength}_{i,i,k}- {Strength}_{i,i,0}$$

Negative values of $\beta_{intra, i,i,k}$ indicate that the effects of test species *i* on itself were more negative on soil conditioned by species *k* than on non-conditioned soil. In other words, the strength of intraspecific competition was more negative on soil conditioned by species *k* than on non-conditioned soil.

After calculating $\beta_{inter, i,j,k}$ and $\beta_{intra, i,i,k}$, we tested whether soil enemies (one aspect of α) explained the soil legacy effect ($\beta_{inter, i,j,k}$ and $\beta_{intra, i,i,k}$; similar with what we did in the Model.link.1). To do so, we used linear mixed models and included the soil legacy effect as the response variable, and diversities of soil bacteria, fungi or fungal pathogens (or relative abundance of fungal pathogens), origin of test focal species and their interaction as fixed effects. We added another fixed effect to distinguish between $\beta_{inter, i,j,k}$ and $\beta_{intra, i,i,k}$ (i.e. the difference between soil-legacy effects on strength of intra- and interspecific competition). We included family and identity of soil-conditioning species, focal test species and competitor test species. To improve homoscedasticity of residuals, soil-conditioning species, focal test species and competitor test species were allowed to vary their residuals.

Finally, we tested whether microbial community dissimilarity (another aspect of α) explained the soil-legacy effect ($\beta_{inter, i,j,k}$ and $\beta_{intra, i,i,k}$; similar with what we did in the Model.link.2). To do so, we used linear mixed models and included the soil-legacy effect as the response variable, and microbial community dissimilarities between soil-conditioning and focal test species, between soil-conditioning and competitor test species and between focal test and competitor test species as fixed effects. We added another fixed effect to distinguish between $\beta_{inter, i,j,k}$ and $\beta_{intra, i,i,k}$. We included family and identity of soil-conditioning species, focal test species and competitor test species as random effects. To improve homoscedasticity of residuals, soil-conditioning species, focal test species and competitor test species were allowed to vary their residuals. Because three out of ten test species were not included in the soil-conditioning phase, we were not able to calculate the microbial community dissimilarity between them and soil-conditioning species, or between them and other test species. Consequently, this analysis was restricted to a small subset (i.e. 268 out of 600 sets), and is less comparable with the results shown in Fig. 3c.

#### S6.1 Effects of diversity of soil bacteria and fungi

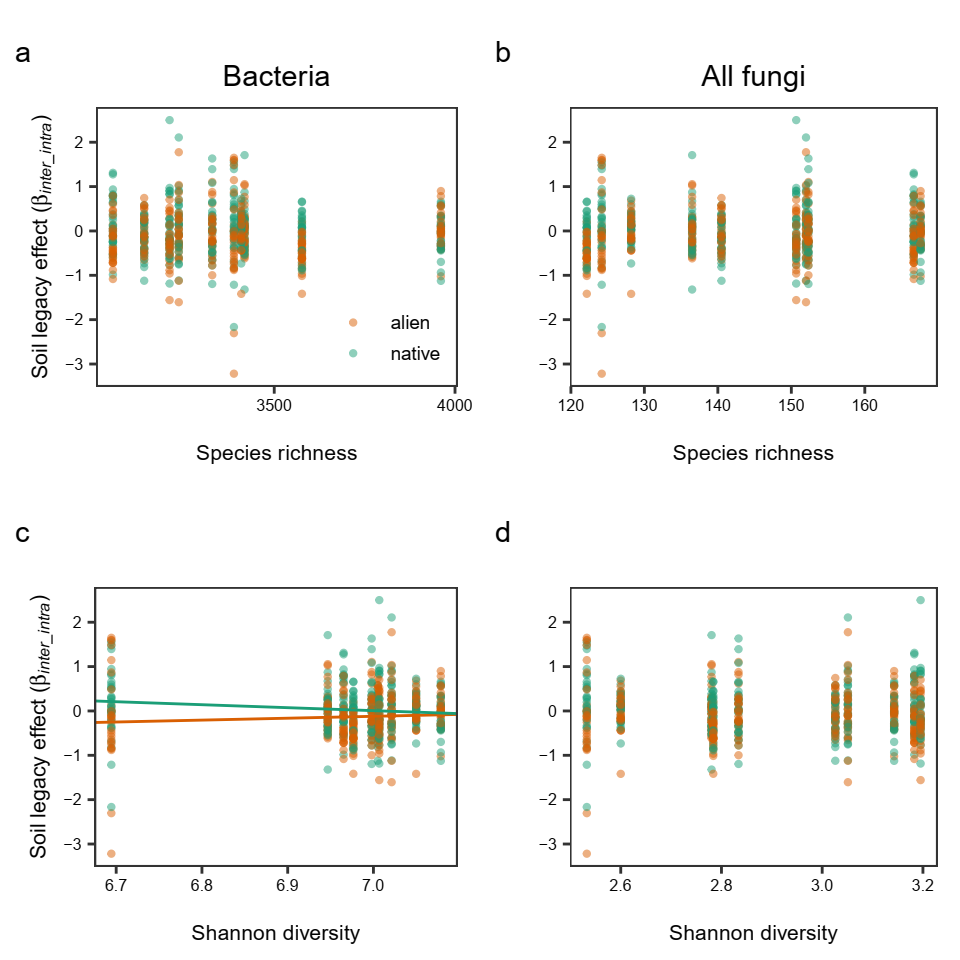

##### Figure S11 Effects of diversity (species richness or Shannon diversity) of bacteria (a, c) and fungi (b, d) on soil-legacy effects.

Green dots represent native test species, and orange dots represent alien test species. Negative values of the soil-legacy effect indicate that the strength of competition became more negative on conditioned soil than on non-conditioned soil.

##### Table S14 Effects of bacterial and fungal diversities **on soil-legacy effects.**

Species richness

|  | Bacteria | |  | All fungi | |
| --- | --- | --- | --- | --- | --- |
|  | χ^2^ | P |  | χ^2^ | P |
| Intra_inter | 0.004 | 0.950 |  | 0.002 | 0.962 |
| Species richness (SR) | 0.006 | 0.940 |  | 1.301 | 0.254 |
| Origin (O) | 0.681 | 0.409 |  | 0.691 | 0.406 |
| O : SR | 0.776 | 0.378 |  | 1.406 | 0.236 |
| **Random effects** | SD |  |  | SD |  |
| Family (soil) | 0.031 |  |  | 0.056 |  |
| Species (soil) | 0.075 |  |  | 0.059 |  |
| Family (focal test) | 0.000 |  |  | 0.000 |  |
| Species (focal test) | 0.228 |  |  | 0.225 |  |
| Family (competitor test) | 0.138 |  |  | 0.137 |  |
| Species (competitor test) | 0.000 |  |  | 0.000 |  |
| Residual | 0.648 |  |  | 0.631 |  |

Shannon diversity

|  | Bacteria | |  | All fungi | |
| --- | --- | --- | --- | --- | --- |
|  | χ^2^ | P |  | χ^2^ | P |
| Intra_inter | 0.004 | 0.952 |  | 0.004 | 0.949 |
| Shannon diversity (Sh) | 0.091 | 0.763 |  | 0.009 | 0.926 |
| Origin (O) | 0.682 | 0.409 |  | 0.681 | 0.409 |
| O : Sh | 4.500 | **0.034*** |  | 0.958 | 0.328 |
| **Random effects** | SD |  |  | SD |  |
| Family (soil) | 0.025 |  |  | 0.026 |  |
| Species (soil) | 0.074 |  |  | 0.077 |  |
| Family (focal test) | 0.000 |  |  | 0.000 |  |
| Species (focal test) | 0.227 |  |  | 0.226 |  |
| Family (competitor test) | 0.137 |  |  | 0.137 |  |
| Species (competitor test) | 0.000 |  |  | 0.000 |  |
| Residual | 0.658 |  |  | 0.632 |  |

#### S6.2 Effect of abundance and diversity of soil fungal pathogens

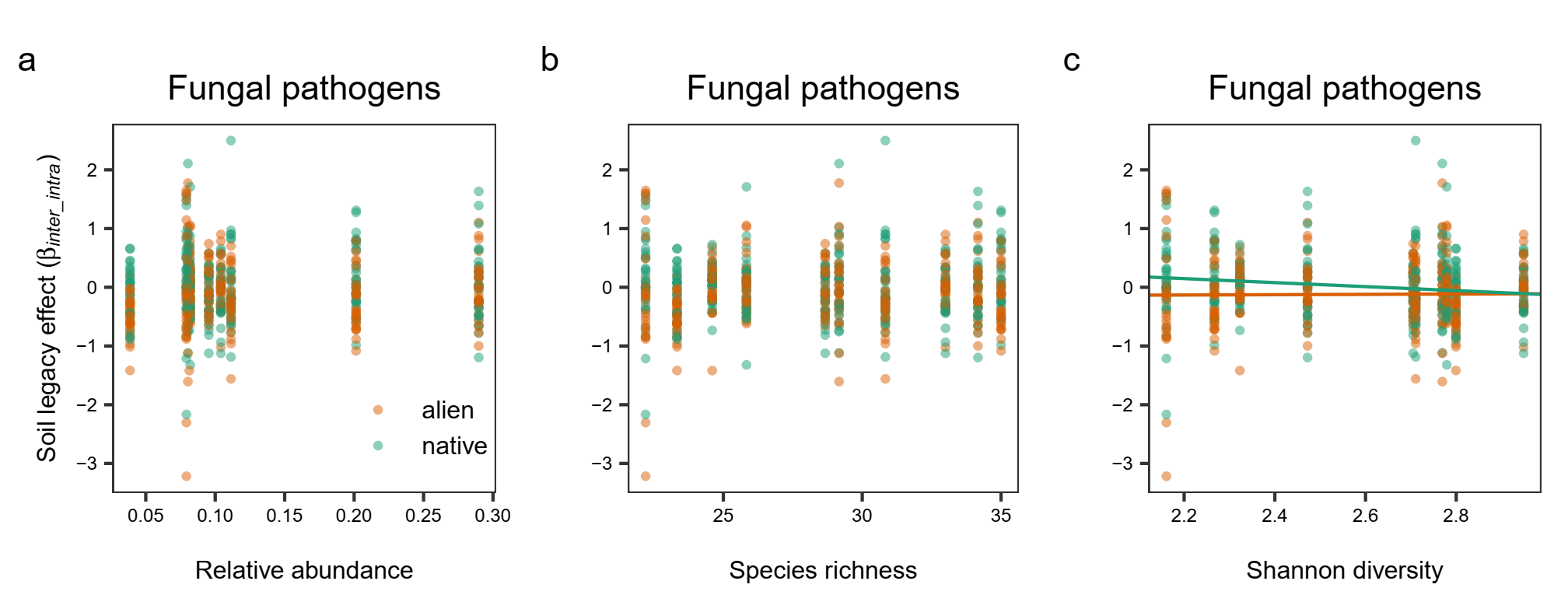

##### Figure S12 Effects of (a) relative abundance, (b) species richness and (c) Shannon diversity of fungal pathogens on soil-legacy effects.

Green dots represent native test species, and orange dots represent alien test species. Negative values of the soil-legacy effect indicate that the strength of competition became more negative on conditioned soil than on non-conditioned soil. Relative abundance was logit-transformed.

##### Table S15 Effect of **fungal pathogens’** abundance and **diversity on soil legacy.**

Relative abundance

|  | Fungal pathogens | |
| --- | --- | --- |
|  | χ^2^ | P |
| Intra_inter^a^ | 0.004 | 0.950 |
| Abundance | 0.173 | 0.677 |
| Origin (O) | 0.683 | 0.409 |
| O : Abundance | 0.427 | 0.514 |
| **Random effects** | SD |  |
| Family (soil) | 0.026 |  |
| Species (soil) | 0.077 |  |
| Family (focal test) | 0.000 |  |
| Species (focal test) | 0.226 |  |
| Family (competitor test) | 0.137 |  |
| Species (competitor test) | 0.000 |  |
| Residual | 0.632 |  |

^a^The difference between changes in strength of intraspecific competition and changes in strength of interspecific competition

Species richness

|  | Fungal pathogens | |
| --- | --- | --- |
|  | χ^2^ | P |
| Intra_inter | 0.003 | 0.955 |
| Species richness (SR) | 0.410 | 0.522 |
| Origin (O) | 0.686 | 0.408 |
| O : SR | 1.800 | 0.180 |
| **Random effects** | SD |  |
| Family (soil) | 0.026 |  |
| Species (soil) | 0.077 |  |
| Family (focal test) | 0.000 |  |
| Species (focal test) | 0.226 |  |
| Family (competitor test) | 0.137 |  |
| Species (competitor test) | 0.000 |  |
| Residual | 0.632 |  |

Shannon diversity

|  | Fungal pathogens | |
| --- | --- | --- |
|  | χ^2^ | P |
| Intra_inter | 0.005 | 0.941 |
| Shannon diversity (Sh) | 0.397 | 0.529 |
| Origin (O) | 0.681 | 0.409 |
| O : Sh | 7.108 | **0.008*** |
| **Random effects** | SD |  |
| Family (soil) | 0.026 |  |
| Species (soil) | 0.077 |  |
| Family (focal test) | 0.000 |  |
| Species (focal test) | 0.226 |  |
| Family (competitor test) | 0.137 |  |
| Species (competitor test) | 0.000 |  |
| Residual | 0.632 |  |

#### S6.3 Effect of soil community dissimilarity on soil-legacy effects

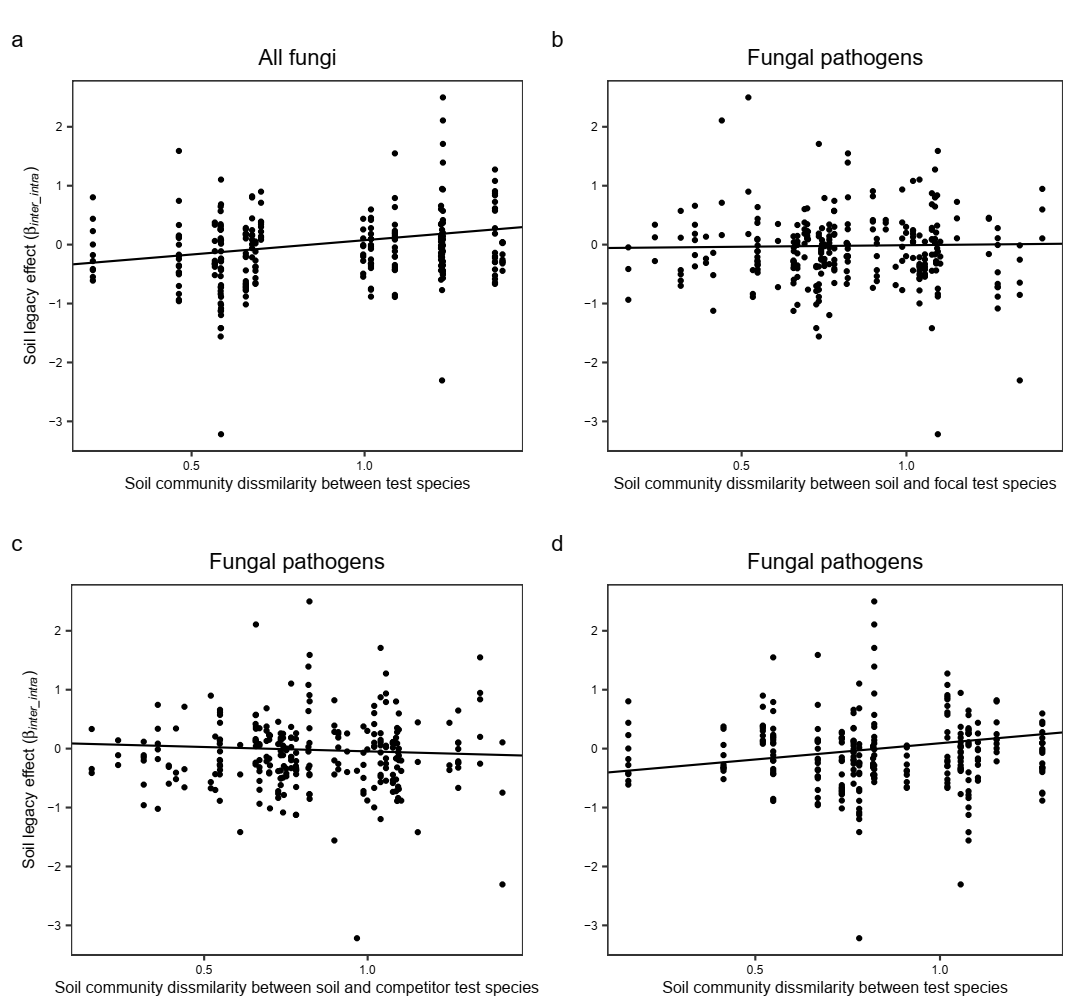

##### **Figure S13 Effects of soil community dissimilarity on soil-legacy effects.** a, dissimilarity of fungal communities between focal test and competitor test species; b, dissimilarity of fungal pathogen communities between soil-conditioning and focal test species; c, dissimilarity of fungal pathogen communities between soil-conditioning and competitor test species; d, dissimilarity of fungal pathogen communities between focal test and competitor test species. Negative values of the soil-legacy effect indicate that the strength of competition became more negative on conditioned soil than on non-conditioned soil. Soil-community dissimilarity was logit-transformed. Only significant effects were plotted.

##### Table S16 Effects of soil community dissimilarity on soil-legacy effects.

|  | Bacteria | |  | All fungi | |  | Fungal pathogen | |  | Fungal endophyte | |
| --- | --- | --- | --- | --- | --- | --- | --- | --- | --- | --- | --- |
|  | χ^2^ | P |  | χ^2^ | P |  | χ^2^ | P |  | χ^2^ | P |
| Intra_inter^a^ | 0.047 | 0.829 |  | 0.578 | 0.447 |  | 0.052 | 0.820 |  | 0.020 | 0.888 |
| Logit(dissimilarity)^b^ | 3.104 | *0.078* |  | 0.262 | 0.609 |  | 3.896 | **0.048*** |  | 1.301 | 0.254 |
| Logit(dissimilarity)^c^ | 2.180 | 0.140 |  | 2.953 | *0.086* |  | 6.462 | **0.011*** |  | 0.554 | 0.457 |
| Logit(dissimilarity)^d^ | 2.094 | 0.148 |  | 15.495 | **0.000*** |  | 5.434 | **0.020*** |  | 0.066 | 0.797 |
| **Random effects** | SD |  |  | SD |  |  | SD |  |  | SD |  |
| Family (soil) | 0.073 |  |  | 0.073 |  |  | 0.081 |  |  | 0.000 |  |
| Species (soil) | 0.000 |  |  | 0.000 |  |  | 0.000 |  |  | 0.000 |  |
| Family (focal test) | 0.000 |  |  | 0.001 |  |  | 0.000 |  |  | 0.000 |  |
| Species (focal test) | 0.329 |  |  | 0.375 |  |  | 0.359 |  |  | 0.300 |  |
| Family (competitor test) | 0.128 |  |  | 0.142 |  |  | 0.159 |  |  | 0.144 |  |
| Species (competitor test) | 0.063 |  |  | 0.000 |  |  | 0.073 |  |  | 0.059 |  |
| Residual | 0.329 |  |  | 0.329 |  |  | 0.330 |  |  | 0.485 |  |

^a^ The difference between changes in strength of intraspecific competition and changes in strength of interspecific competition

^b^ Dissimilarity in soil-community composition between soil-conditioning and focal test species

^c^ Dissimilarity in soil-community composition between soil-conditioning and competitor test species

^d^ Dissimilarity in soil-community composition between focal and competitor test species

### Supplement S7 Phylogenetic relatedness and soil community dissimilarity

We constructed a phylogenetic tree for our soil-conditioning species by pruning the phylogenetic tree constructed by Smith and Brown ^1^, which is hitherto the most comprehensive tree for seed plants. As *Leontodon autumnalis* is a synonym of *Scorzoneroides autumnalis*, we used the latter for matching.

We used the Mantel test to test for significance of the correlation between phylogenetic distance and soil-community dissimilarity. Then, we used the Kruskal-Wallis test to test whether phylogenetic relatedness is affected by the type of species combination, that is, between two aliens, between an alien and a native, and between two natives.

Although phylogenetic distance correlated positively with dissimilarity of bacterial and fungal endophyte communities (Table S10), phylogenetic distance was not related to the type of species combination (df = 2, χ^2^ = 1.05, P = 0.592).

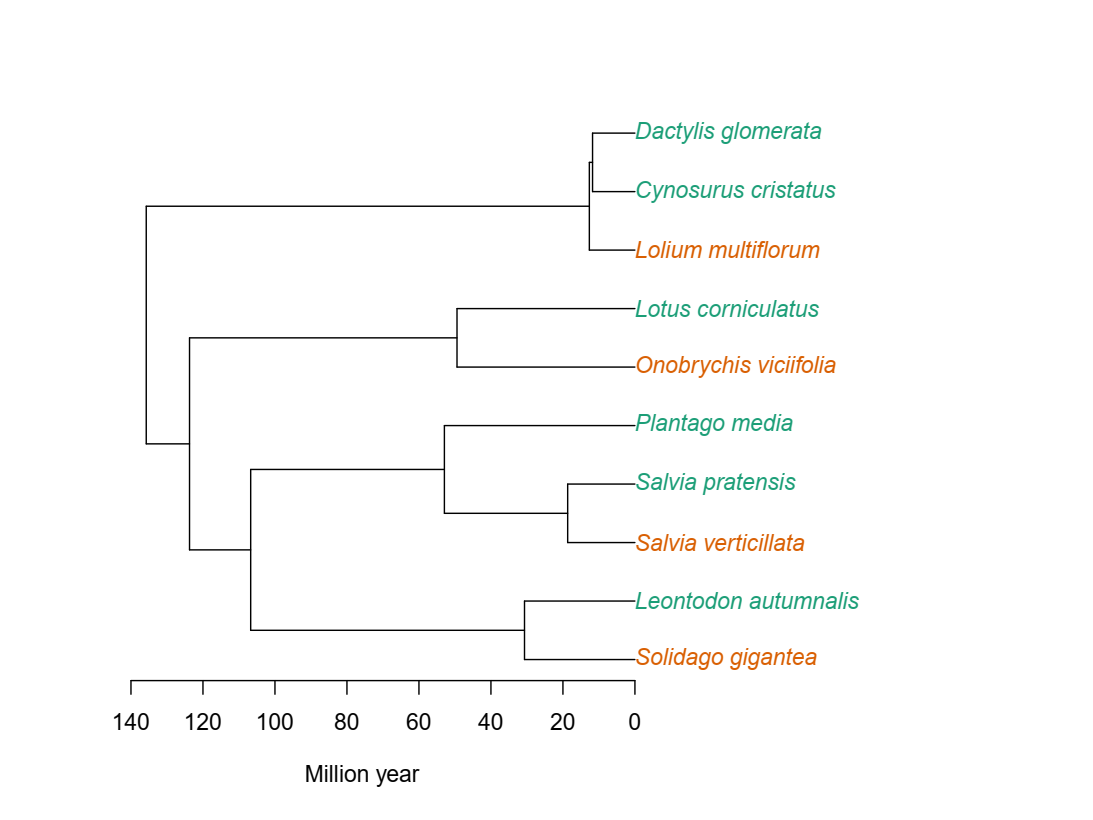

##### Figure S14 Phylogenetic tree of the ten soil-conditioning species.

Green indicates native species, and orange indicates alien species.

##### Table S17 Relationship between phylogenetic distance and soil-community dissimilarity

Significant effects (P < 0.05) are in bold and marked with asterisks.

| Bacteria | |  | All fungi | |  | Fungal pathogens | |  | Fungal endophytes | |
| --- | --- | --- | --- | --- | --- | --- | --- | --- | --- | --- |
| r | P |  | r | P |  | r | P |  | r | P |
| 0.354 | **0.017*** |  | -0.086 | 0.754 |  | 0.070 | 0.263 |  | 0.197 | **0.040*** |

The significance of the correlation between phylogenetic distance and soil community dissimilarity was tested with a Mantel test.
